# SARS-CoV-2 genome-wide mapping of CD8 T cell recognition reveals strong immunodominance and substantial CD8 T cell activation in COVID-19 patients

**DOI:** 10.1101/2020.10.19.344911

**Authors:** Sunil Kumar Saini, Ditte Stampe Hersby, Tripti Tamhane, Helle Rus Povlsen, Susana Patricia Amaya Hernandez, Morten Nielsen, Anne Ortved Gang, Sine Reker Hadrup

**Author notes:** These authors contributed equally to this work (Ditte Stampe Hersby, Tripti Tamhane).

## Abstract

To understand the CD8^+^ T cell immunity related to viral protection and disease severity in COVID-19, we evaluated the complete SARS-CoV-2 genome (3141 MHC-I binding peptides) to identify immunogenic T cell epitopes, and determine the level of CD8^+^ T cell involvement using DNA-barcoded peptide-major histocompatibility complex (pMHC) multimers. COVID-19 patients showed strong T cell responses, with up to 25% of all CD8^+^ lymphocytes specific to SARS-CoV-2-derived immunodominant epitopes, derived from ORF1 (open reading frame 1), ORF3, and Nucleocapsid (N) protein. A strong signature of T cell activation was observed in COVID-19 patients, while no T cell activation was seen in the ‘non-exposed’ and ‘high exposure risk’ healthy donors. Interestingly, patients with severe disease displayed the largest T cell populations with a strong activation profile. These results will have important implications for understanding the T cell immunity to SARS-CoV-2 infection, and how T cell immunity might influence disease development.

## Introduction

The COVID-19 (Coronavirus disease 2019) pandemic caused by the highly infectious SARS-CoV-2 (severe acute respiratory syndrome coronavirus 2) has challenged public health at an unprecedented scale, causing the death of more than one million people so far (World Health Organization). The T-cell of the immune system is the main cell type responsible for the control and elimination of viral infections; CD8^+^ T cells are critical for the clearance of viral-infected cells, whereas CD4^+^ T cells are critical for supporting both the CD8^+^ T cell response efficacy and the generation of specific antibodies. Characteristics from the ongoing pandemic suggest that T cell recognition will be critical to mediate long-term protection against SARS-CoV-2 (Cañete and Vinuesa, 2020), as the antibody-mediated response seems to decline in convalescent patients in 3-month follow-up evaluation (Seow et al., 2020; Vabret, 2020). Furthermore, studies of the closely related SARS-CoV infection shows persistent memory CD8^+^ T cell responses even after 11 years in SARS recovered patients without B cell responses (Peng et al., 2006; Tang et al., 2011), emphasizing the potential role of CD8^+^ memory T cells in long-term protection from coronaviruses.

Several recent studies have reported robust T cell immunity in SARS-CoV-2 infected patients (Ni et al., 2020; Peng et al., 2020; Weiskopf et al., 2020), and unexposed healthy individuals also showed functional T cell reactivity restricted to SARS-CoV-2 (Le Bert et al., 2020; Grifoni et al., 2020; Meckiff et al., 2020; Nelde et al., 2020; Ni et al., 2020; Sekine et al., 2020). The T cell cross-reactivity is hypothesized to derive from routine exposure to common cold coronaviruses (HCoV) (HCoV-OC43, HCoV-HKU1, HCoV-NL63 and HCoV-229E) that are widely circulated in the population with 90% of the human population being seropositive for these viruses (Braun et al., 2020; Kissler et al., 2020), which share sequence homology with the SARS-CoV-2 genome (Mateus et al., 2020; Stervbo et al., 2020).

SARS-CoV-2 infection might result in mild to severe disease (including death), but also a large number of asymptomatic infections are described (Havers et al., 2020; Huang et al., 2020; Oran and Topol, 2020). The presence of pre-existing T cell immunity, represented by cross-reactive T cells, could have marked implications for how individuals respond to SARS-CoV-2 infection. However, their biological role upon encounter with SARS-CoV-2 infection remains unclear, and their contribution to disease protection needs to be determined. Furthermore, in severe clinical disease, cytokine release syndrome is reported, and might, in some cases, be dampened to immunosuppressive medication or anti-IL6 antibody-therapy (Moore and June, 2020; Zhang et al., 2020). Such clinical characteristics point to a potential uncontrolled immune response with the involvement of strong T cell activation.

T cells are activated by a specific interaction between the T cell receptor (TCR) and peptide-antigen presented by major histocompatibility complex (MHC) molecules on the surface of virus-infected cells. Although SARS-CoV-2-specific immunity has been reported both in the context of COVID-19 infection and pre-existing T cells, the exact antigens (minimal peptide epitope) of the viral genome associated with this immunity in COVID-19-infected patients is not fully known. Using our large-scale T cell detection technology based on DNA-barcoded peptide-MHC multimers (Bentzen et al., 2016), we have mapped T cell recognition throughout the complete SARS-CoV-2 genome, and identified the exact epitopes recognized by SARS-CoV-2-specific CD8^+^ T cells. Furthermore, we have evaluated the phenotype characteristics of T cells recognizing these epitopes in COVID-19 patients and in healthy donors.

## Results

### SARS-CoV-2-specific CD8 T cell immunity is represented by strong immunodominant epitopes

To reveal the full spectrum of T cell immunity in COVID-19 infection, we used a complete SARS-CoV-2 genome sequence (Wu et al., 2020) to identify immunogenic minimal epitopes recognized by CD8^+^ T cells. Using NetMHCpan 4.1 (Reynisson et al., 2020), we selected 2204 potential HLA binding peptides (9–11 amino acids) for experimental evaluation. These peptides were predicted to bind one or more of ten prevalent and functionally diverse MHC class I (MHC-I) molecules representing HLA-A (A01:01, A02:01, A03:01, and A24:02), -B (B07:02, B08:01, B15:01), and -C (C06:02, C07:01, and C07:02) loci, leading to a total 3141 peptide-MHC specificities for experimental evaluation (Figure 1A, Supplementary Table 1, Supplementary Figure 1A). T cell reactivity towards these peptides was analyzed in 18 COVID-19-infected patients (Supplementary Table 2), with a mean HLA coverage of 3.1 and evaluation of an average of 972 peptides per patient (Supplementary Figure 1B) by our high-throughput T cell detection technology using DNA-barcoded peptide-MHC (pMHC) multimers (Bentzen et al., 2016). Briefly, each pMHC complex is multimerized on a PE-(Phycoerythrin) or APC-(Allophycocyanin) labeled dextran backbone and tagged with a unique DNA-barcode. DNA-barcoded pMHC multimers are then pooled to generate an HLA matching patient-tailored pMHC multimer panel, which is incubated with patient-derived PBMCs (peripheral blood mononuclear cells), and multimers bound to CD8^+^ T cells are sorted and sequenced to identify T cell recognition towards the probed pMHC complexes. For phenotype characterization of SARS-CoV-2-specific CD8^+^ T cells, we combined pMHC multimer analysis with a 13-parameter antibody panel (Supplementary Table 3), and for comparative evaluation, we furthermore included 39 T cell epitopes from common viruses; cytomegalovirus (CMV), Epstein-Barr virus (EBV), and influenza (Flu) virus (CEF, Supplementary Table 4) (Figure 1B).

**Figure 1.**
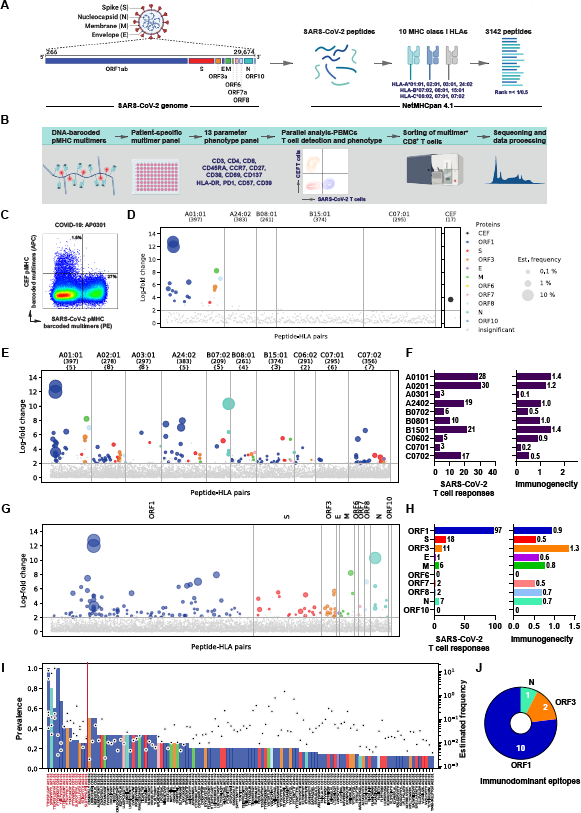
CD8^+^ T cell epitope mapping in SARS-CoV-2. (**A**) Schematic representation of the complete SARS-CoV-2 genome used for the identification of 3141 peptides with predicted binding rank (NetMHCpan 4.1) of ≤ 0.5 (ORF1 protein) and ≤ 1 (all remaining proteins) for ten prevalent HLA-A, B, and C molecules. (**B**) Experimental pipeline to analyze for T cell recognition towards the 3141 SARS-CoV-2-derived HLA-binding peptides in PBMCs using DNA-barcoded peptide-MHC (pMHC) multimers. A 13-antibody panel was used for phenotype analysis of pMHC multimer positive CD8^+^ T cells. pMHC multimers binding CD8^+^ T cells were sorted based on PE (SARS-CoV-2-specific) or APC (CEF-specific) signal, and sequenced to identify antigen-specific CD8^+^ T cells. (**C**) Representative flow cytometry plot of CD8^+^ T cells from a COVID patient stained with pMHC multimer panel showing SARS-CoV-2 (PE) and CEF (APC) multimer^+^ T cells that were sorted for DNA-barcode analysis to identify epitopes recognition. **D**) CD8^+^ T cell recognition to individual epitopes were identified based on the enrichment of DNA barcodes associated with each of the tested peptide specificities (LogFc>2 and p < 0.001, *barracoda*). Significant T cell recognition of individual peptide sequences are colored based on their protein of origin and segregated based on their HLA-specificity. The size of the dot represents the estimated frequency of pMHC multimer positive CD8^+^ T cell populations for each of the recognized epitopes. The black dots show CD8^+^ T cells reactive to one of the CEF peptides (here, CMV pp65; YSEHPTFTSQY-HLA-A01:01). All peptide sequences with no significant enrichments are shown as gray dots. (**E**) Summary of all the T cell recognition to SARS-CoV-2-derived peptides identified in the 18 analyzed COVID-19-patients. In parentheses, number of peptides tested for each HLA (upper row) and the number of patients analyzed for each HLA pool (lower row). Each dot represents one peptide-HLA combination per patient. The dot size represents the estimated frequency of each pMHC specific CD8^+^ T cell response (as in panel C), and are colored according to their origin of protein, similar to shown in panel A. (**F**) Bar plots summarize the number of HLA-specific epitopes and the HLA-restricted immunogenicity (given as T cell recognition) in the analyzed patient cohort. Immunogenicity represents the fraction of T cell recognized peptides out of the total number of peptides analyzed for a given HLA-restriction across the HLA-matching donors (given as %). (**G**) Similar to E, a summary of T cell responses separated based on the protein of origin. (**H**) Bar plots show the number of epitopes derived from each of the SARS-CoV-2 protein and their immunogenicity score (%). (**I**) The prevalence of T cell recognition towards the individual epitopes detected in COVID-19 patients (left Y-axis) and the estimated frequency for each of the T cell populations detected (right Y-axis). Dotted line indicates the epitopes determined as immunodominant. Bars are colored according to their protein of origin, similar as shown in panel A. (**J**) Pie chart of immunodominant epitopes distributed according to their protein of origin.

We found broad and strong SARS-CoV-2-specific CD8^+^ T cell responses in COVID-19 patients, contributing up to 27% of the total CD8^+^ T cells (Figure 1C). A few epitopes showed characteristics of immunodominance, and raised T cell responses up to 25% of total CD8^+^ T cells against two overlapping epitopes; HLA-A01:01 TTDPSFLGRY (11%), and TTDPSFLGRYM (14%), in some patient-specific T cell repertoires (Figure 1D, Supplementary Figure 2, and Supplementary Table 5). We validated the presence of SARS-CoV-2 CD8^+^ T cells identified with our pMHC multimer-based T cell detection technology using functional analysis in selected patient samples (Supplementary Figure 3).

In total, we identified T cell responses to 142 pMHC complexes corresponding to 122 unique T cell epitopes across the ten analyzed HLAs (Figure 1E). HLA-A01:01, A02:01, and B15:01 dominated in terms of the total number of identified epitopes as well as the immunogenicity score (i.e., the number of T cell responses normalized to the number of probing pMHC multimers and the number of patients analyzed) (Figure 1F). HLA-A03:01 and C07:01 specific peptides showed the least T cell reactivity (three epitopes each) despite being analyzed in nine and six patients, respectively (Figure 1E). Most of the immunogenic epitopes were mapped to the ORF1 protein, followed by S and ORF3 proteins (Figure 1G and H, and Supplementary Table 5). Given the size difference of the viral proteins, the ‘immunogenicity score’ was used to evaluate their relative contribution to T cell recognition. Based on such evaluation, we observe that peptides derived from ORF3 displayed the highest relative immunogenicity (in terms of T cell recognition), followed by ORF1 protein (Figure 1H). Of the 122 epitopes recognized by T cells in the patient cohort, 13 were determined as ‘immunodominant’ based on the presence of T cell recognition in two or more patients, and prevalence of >25% in the tested samples with the given HLA molecule (Figure 1I). Among these, a very robust HLA associated immunodominance was observed for two of the epitopes: HLA-A01:01-TTDPSFLGRY-specific (and its variant peptides TTDPSFLGRYM and HTTDPSFLGRY), with specific T cells detected in all five analyzed patients (estimated frequency reaching up to 25% of total CD8^+^ T cells); and HLA-B07:02-SPRWYFYYL, with specific T cells observed in four of the five patients evaluated (estimated frequency up to 10%) (Figure 1I). Surprisingly, in our patient cohort, none of the immunodominant epitopes were derived from the S protein, despite this being the second-largest protein (Figure 1J).

In summary, we report SARS-CoV-2-specific CD8^+^ T cell immunity towards several epitopes and the recurrent finding of T cell recognition towards a few immunodominant epitopes. The ORF1 protein contributes the most to T cell recognition of SARS-CoV-2, but is also by far the largest group of proteins. When protein size is taken into account, ORF3 and ORF1 contributed the most for both immunogenic and immunodominant epitopes.

### Large-scale reactivity towards SARS-CoV-2-derived peptides in healthy individuals

Next, we analyzed healthy individuals for T cell recognition against all 3141 SARS-CoV-2-derived peptides. We selected two healthy donor cohorts; representing COVID-19 unexposed healthy individuals (HD-1; n=18, PBMCs collected before COVID-19 outbreak), and healthcare staff at high-risk of SARS-CoV-2 exposure but not tested COVID-19 positive (HD2; n=20, PBMCs collected during COVID-19 outbreak). CD8^+^ T cells of COVID-19 unexposed healthy individuals showed large-scale T cell recognition towards SARS-CoV-2-derived peptides from across the whole viral genome (Figure 2A, Supplementary Figure 4, and Supplementary Table 6a). Cumulatively, 214 SARS-CoV-2-derived peptides were recognized by T cells in 16 out of the 18 analyzed samples. Despite such broad T cell-recognition, most of the T cell populations were of low-frequency, never exceeding 1% of total CD8^+^ T cells. The high-risk COVID-19 healthy cohort showed similar T cell recognition towards 178 SARS-CoV-2 epitopes (Figure 2B, Supplementary Figure 4, and Supplementary Table 6b). SARS-CoV-2 recognizing T cell populations in this cohort were also of low frequency (less than 1% of total CD8^+^ T cells) and were identified in 15 of the 20 donors. Interestingly, while T cell recognition was seen at a substantial level in both the healthy donor cohorts; the staining-index of the pMHC multimer binding in healthy donors was significantly lower than similarly observed in patients (Figure 2C). This might indicate a lower TCR avidity to the probed pMHC in healthy individual compared to COVID-19 patients, and could be due to potential cross-reactivity from existing T cell populations raised against other coronaviruses (such as common cold viruses HCoV-HKU1, HCoV-229E, HCoV-NL63, and HCoV-OC43) that share some level of sequence homology with SARS-CoV-2, as suggested in recent reports (Le Bert et al., 2020; Braun et al., 2020; Mateus et al., 2020).

**Figure 2.**
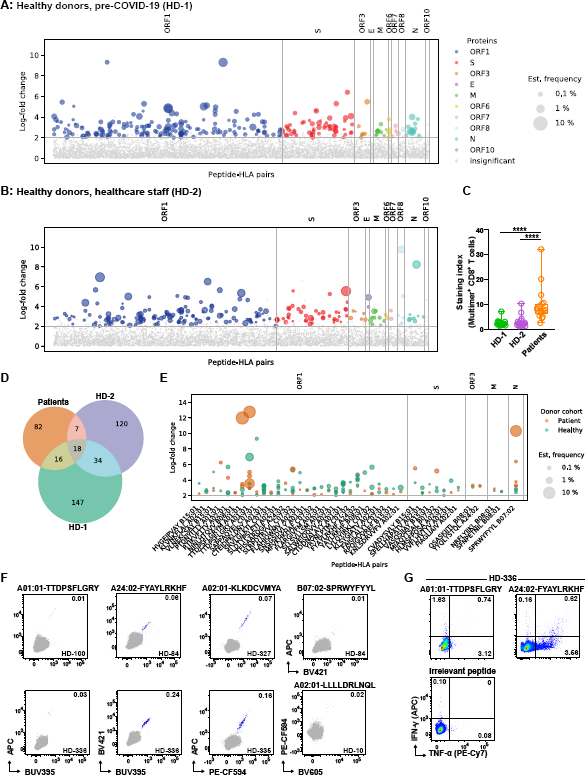
Large-scale reactivity towards SARS-CoV-2-derived peptides in healthy individuals. (**A**) CD8^+^ T cell recognition to individual SARS-CoV-2-derived peptides in the pre-COVID-19 healthy donor cohort (n=18) identified based on the enrichment of DNA barcodes associated with each of the tested peptide specificities (LogFc>2 and p < 0.001, *barracoda*). Significant T cell recognition of individual peptide sequences are colored and segregated based on their protein of origin. The size of the dot represents the estimated frequency of pMHC multimer positive CD8^+^ T cell populations for each of the recognized peptides (**B**) Similar to A, but displays the high-exposure risk healthy donor cohort (n=20). (**C**) Staining index of CD8^+^ T cells binding SARS-CoV-2-specific pMHC multimers in the three evaluated cohorts. The staining index is calculated as; ((mean fluorescence intensity (MFI) of multimer^+^ cells – MFI of multimer^-^ cells)/2 × standard deviation (SD) of multimer^-^ cells)). Mann-Whitney test; p < 0.0001 (patient vs. HD-1), p < 0.0001 (patient vs. HD-2), and p = 0.86 (HD-1 vs. HD-2); n = 18 (patient), n = 18 (HD-1), and n = 20 (HD-2). (**D**) Venn diagram illustrating the overlap of T cell recognition towards SARS-CoV-2-derived peptides in COVID-19 patient and healthy donor cohorts. (**E**) Comparison of estimated T cell frequency (size of the circle) towards the 41 SARS-CoV-2-derived peptides recognized both in patients and healthy donors. (**F**) Flow cytometry dot plots showing *in-vitro* expanded T cells from healthy donors recognizing SARS-CoV-2 derived epitopes, detected by combinatorial tetramer staining. T cell binding to each pMHC specificity is detected using pMHC tetramers prepared in a two-color combination (blue dots), gray dots show tetramer negative T cells, and the number on the plots shows the frequency (%) of tetramer^+^ of the CD8^+^ T cells. (**G**) Intracellular cytokine staining (TNFα and INFγ) of SARS-CoV-2 expanded T cell populations from healthy donor (HD-336, panel F) following antigen stimulation. The numbers on the plot indicate the % frequency of CD8^+^ T cells double or single positive for the analyzed cytokines. HLA-A01:01 restricted irrelevant peptide was used as a negative control. The gating strategy of the flow cytometry analysis is shown in **Supplementary Figure 5**.

Forty-one of the COVID-19 immunogenic peptides, including eight of the immunodominant peptides, identified in the patient cohort were also recognized by T cells of healthy donors (Figure 2D), this includes the two most frequently observed epitopes of SARS-CoV-2: HLA-A01:01-TTDPSFLRGY and HLA-B07:02-SPRWYFYYL (Figure 2E). For further validation, we expanded T cells *in-vitro* from several COVID-19 unexposed healthy donors and measured T cell binding using fluorophore-labeled pMHC tetramers. Based on *in-vitro* peptide-driven expansion pMHC tetramer binding T cell populations were verified in multiple donors for SARS-CoV-2-derived peptides, including immunodominant epitopes across four HLAs (A01:01-TTDPSFLRGY, A02:01-LLLLDRLNQL, A02:01-KLKDCVMYA, A24:02-FYAYLRKHF, and B07:02-SPRWYFYYL) (Figure 2F). Although these T cell responses were of low magnitude, a functional response (measured by IFN-γ and TNF-α) was observed in these *in-vitro* expanded T cell cultures when re-stimulated with individual peptide epitopes (Figure 2G) or epitope pools (Supplementary Figure 5B).

Altogether, we show a full spectrum of functionally validated T cell recognition towards SARS-CoV-2-derived peptides in healthy donors; this is detected at low frequency and seems to have low-avidity interaction (as determined based on the staining index of the pMHC multimer interaction).

### Strong and distinct activation profile exhibited by SARS-CoV-2-specific T cells in COVID-19 patients

Since our study design integrated T cell phenotype characterization in combination with the SARS-CoV-2-specific T cell identification, we evaluated and compared phenotypic characteristics of the SARS-CoV-2 reactive T cell populations in COVID-19-infected patients and healthy donors. This also allowed us to evaluate whether the multimer-specific T cell profile of the high-risk COVID-19 healthy cohort (HD-2) has any distinct features compared to the unexposed cohort (HD-1). Dimensionality reduction visualization of SARS-CoV-2 multimer positive T cells using UMAP (Uniform Manifold Approximation and Projection) showed distinct clustering for the patient cohort compared to the two healthy donor cohorts (Figure 3A). Patient-derived SARS-CoV-2 multimer-positive T cells were distinguished with higher expression of activation markers CD38, CD69, CD39, HLA-DR, CD57, and reduced expression of CD8 and CD27 (Figure 3B). These features were found unique to SARS-CoV-2-specific T cells, as no difference was observed between the three cohorts in similar analysis for CEF-specific multimer positive T cells (Supplementary Figures 6A and B). Additionally, SARS-CoV-2 reactive T cells in patients and healthy donor cohorts showed a similar distribution of memory subsets, effector memory (EM) CCR7^-^ CD45RA^-^; central memory (CM), CCR7^+^ CD45RA^-^; and terminally differentiated effector memory (TEMRA), CCR7^-^CD45RA+; and naïve (CCR7+, CD45RA+) phenotype (Supplementary Figures 7A and B). However, SARS-CoV-2-specific T cells of the patient cohort showed unique clustering (UMAP) of EM and TEMRA and these two subsets particularly showed strong expression of the T cell activation markers (Supplementary Figures 7C and D) compared to both healthy donor cohorts.

**Figure 3.**
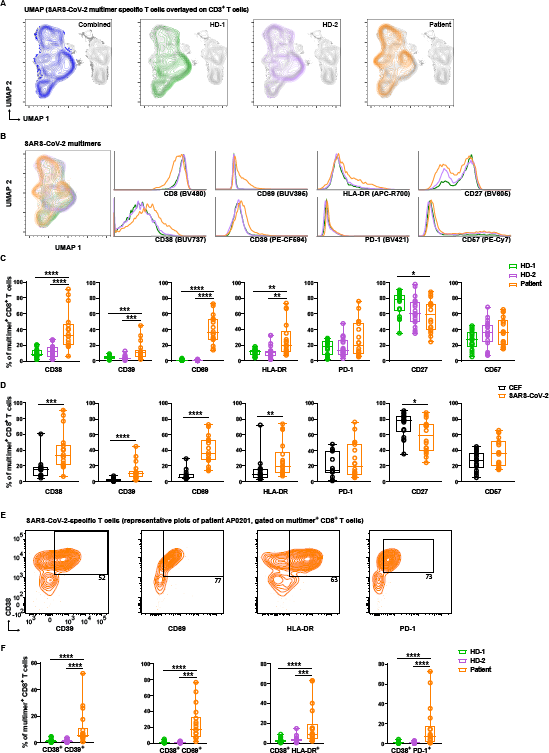
Strong and distinct activation profile of SARS-CoV-2-specific T cells in COVID-19 patients. (**A**) UMAP summarizing clustering of SARS-CoV-2 pMHC multimer binding CD8^+^ T cells overlaid on CD3^+^ cells showing combined (blue) and individual distribution of patient (orange) and two healthy donor cohorts (green and purple). (**B**) UMAP overlay of SARS-CoV-2 pMHC multimer binding CD8^+^ T cells in the three cohorts and histograms showing comparative expression of individual phenotype markers (n = 18 for each cohort). (C) Box plots comparing frequencies of SARS-CoV-2 pMHC multimer binding CD8^+^ T cells carrying the individual of phenotype markers indicated, in the 3 different cohorts (n = 18 for each cohort). Each dot represent one sample. Frequencies were quantified from flow cytometry data processed using the gating strategy applied in **Supplementary Figure 9**. P-values for hypothesis (Mann-Whitney) test; CD38 (p < 0.0001 HD-1 vs. patient, p < 0.0001 HD-2 vs. patient), CD39 (p = 0.0001 HD-1 vs. patient, p = 0.001 HD-2 vs. patient), CD69 (p < 0.0001 HD-1 vs. Patient, p < 0.0001 HD-2 vs. patient), HLA-DR (p = 0.002 HD-1 vs. patient, p = 0.002 HD-2 vs. patient), and CD27 (p = 0.01 HD-1 vs. patient). (**D**) Box plots comparing SARS-CoV-2 pMHC multimer^+^ (n = 18) and CEF pMHC multimer^+^ (n = 14) CD8^+^ T cells identified in the COVID-19 patient cohort for the expression of phenotype markers. Each dot represent one sample. P-values for hypothesis (Mann-Whitney) test; p = 0.0002 (CD38), p < 0.0002 (CD39), p < 0.000 (CD69), p = 0.009 (HLA-DR), p = 0.01 (CD27). (**E**) Representative flow cytometry plots of SARS-CoV-2 pMHC multimer^+^ CD8^+^ T cells of a COVID-19 patient sample showing expression of activation markers (CD39, CD69, and HLA-DR) and PD-1 in combination with CD38. Numbers on the plots show the frequency the gated population. (**F**) Quantification of SARS-CoV-2 pMHC multimer^+^ CD8^+^ T cells expressing the phenotype profile shown in the representative plots (upper row) comparing patient and healthy donor cohorts (n = 18 for all three cohorts). P-values for hypothesis (Mann-Whitney) test; CD38^+^ CD39^+^ (p < 0.0001 HD-1 vs. patient, p < 0.0001 HD-2 vs. patient), CD38^+^ CD69^+^ (p < 0.0001 HD-1 vs. patient, p < 0.001 HD-2 vs. patient), CD38^+^ PD-1^+^ (p < 0.0001 HD-1 vs. patient, p < 0.0001 HD-2 vs. patient), CD38^+^ HLA-DR^+^ (p < 0.0001 HD-1 vs. patient, p = 0.0001 HD-2 vs. patient).

Phenotype analysis demonstrated a highly activated state of SARS-CoV-2-specific T cells, with a significantly higher fraction of such T cells expressing the inflammation marker (CD38) and early-stage activation markers (CD39, CD69, and HLA-DR), and showed a late-differentiated effector memory profile (reduced CD27) together with increased CD57 expression (not significant) compared to the two healthy donor cohorts (Figure 3C). We did not observe activation of SARS-CoV-2 specific multimer-positive T cells in the high-risk COVID-19 healthy cohort, except for non-significant trends for reduced CD27 and increased CD57 expression (Figure 3C). Furthermore, the highly activated and differentiated T cell phenotype in the COVID-19 patients was associated with the SARS-CoV-2-specific T cells and not to the CEF-specific T cells detected in this cohort (Figure 3D). To further characterize the SARS-CoV-2-specific T cell phenotype, we compared the expression of the T cell activation markers in combination with inflammatory response marker CD38 on multimer positive CD8^+^ T cells across the three cohorts, which showed significantly enhanced expression of activation molecules (CD39, CD69, and HLA-DR) and PD-1 inhibitory receptor on CD38^+^ T cells only in the patient cohort (Figure 3E and F).

Altogether, these results demonstrate highly active and proliferative SARS-CoV-2-specific T cell responses in COVID-19-infected patients, and distinguishes them from the potential cross-reactive T cell repertoire detected in healthy donors.

### The severity of COVID-19 disease correlates with enhanced activation of SARS-CoV-2-specific CD8 T cells

To dissect the association of T cells with COVID-19 disease severity, we next evaluated the phenotype characteristics of SARS-CoV-2-specific CD8^+^ T cells in the patient cohort related to their requirement for hospital care. Our patient cohort consists of severely diseased patients requiring hospitalization (hospitalized; n = 11) and patients with mild symptoms not requiring hospital care (outpatient; n = 7). Substantially higher frequency of SARS-CoV-2-specific CD8^+^ T cells and increased total number of epitopes were detected in samples from hospitalized patients compared to outpatient samples (Figure 4A). Comparing the SARS-CoV-2 specific T cell population (multimer+) between hospitalized and outpatients for phenotype markers, we observed a clear trend in increased expression of CD38, CD39, CD69, HLA-DR (non-significant), and PD-1 (significant) (Figure 4B). Furthermore, measuring co-expression of immune activation marker CD38 together with CD39, CD69, PD-1, and HLA-DR showed a strong elevation of these combinations of activation markers in hospitalized patients (Figure 4C and D), suggesting a role for SARS-CoV-2 CD8^+^ T cells in severe COVID-19 infection. It should be noted that samples from hospitalized patients might have been collected relatively later after the time-of-infection compared to the outpatient cohort, which could influence the level of T cell reactivity. To solve this potential bias, ongoing studies are addressing (in detail) the kinetics of the T cell response after time-of-infection.

**Figure 4.**
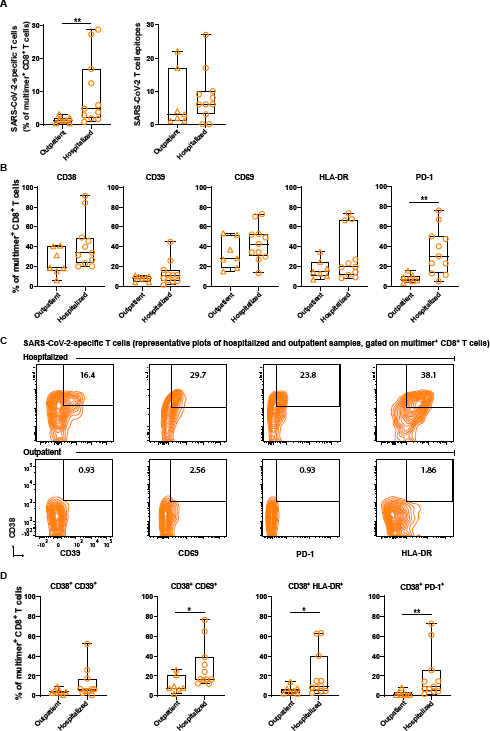
The severity of COVID-19 infection correlates with enhanced activation of SARS-CoV-2 specific CD8^+^ T cells. (**A**) **Left**, Box plots shows frequencies of SARS-CoV-2 multimer^+^ CD8^+^ T cells in outpatient (n = 7) and hospitalized patients (n = 11). P-value (Mann-Whitney test) = 0.008. **Right**, Number of identified SARS-CoV-2 specific T cell epitopes in outpatient (n = 7) and hospitalized (n = 11) patient samples. (**B**), Box plots showing the fraction of SARS-CoV-2 pMHC multimer positive CD8^+^ T cells expressing the indicated phenotype markers in outpatients (n = 7) and hospitalized patients (n = 11). Each dot represents one sample. Frequencies were quantified from flow cytometry data processed using the gating strategy applied in **Supplementary Figure 9**. P-values for hypothesis (Mann-Whitney) test; p = 0.003 (PD-1). (C) Representative flow cytometry plots of SARS-CoV-2 pMHC multimer^+^ CD8^+^ T cells showing expression of activation markers (CD39, CD69, and HLA-DR) and PD-1 in combination with CD38, either in hospitalized (top panel) or outpatient (bottom panel) samples. The numbers on the plots show the frequency of the gated population. (D) Quantification of the SARS-CoV-2 pMHC multimer+ CD8^+^ T cells expressing the phenotype profile shown in the representative plots (top) comparing outpatients (n = 7) and hospitalized patients (n = 11). P-values for hypothesis (Mann-Whitney) testing; p = 0.02 (CD38^+^ CD69+), p = 0.002 (CD38^+^ PD-1+), and p = 0.04 (CD38^+^ HLA-DR^+^).

### A fraction of SARS-CoV-2 epitopes share sequence homology with widely circulating common cold coronaviruses

Pre-existing T cells immunity, in the context of SARS-CoV-2-reactive T cells in COVID-19 unexposed healthy individuals, has been reported by several studies (Le Bert et al., 2020; Braun et al., 2020; Grifoni et al., 2020; Mateus et al., 2020; Nelde et al., 2020), and it has been hypothesized that this is due to the shared sequence homology between the SARS-CoV-2 genome and other common cold coronaviruses (HCoV-OC43, HCoV-HKU1, HCoV-NL63, and HCoV-229E). Having evaluated the full spectrum of minimal epitopes for T cell recognition, we set out to evaluate the sequence homology at the peptide level, and its correlation with the SARS-CoV-2 T cell reactivity observed in the healthy donors. First, we searched for any immunogenic hot-spots across the full SARS-CoV-2 proteome by comparing the number of identified epitopes (in the patient cohort) to the total number of predicted peptides in any given region of the proteins. Generally, the epitopes were spread over the full length of the protein sequences, while clustering in minor groups throughout all regions of the viral proteome (Figure 5A). Regions indicated by asterisk, demonstrates significant enrichment of T cell recognition, relative to the number of MHC-I binding peptides in a given region. Both the C- and N-terminal regions of the ORF1 seem to hold fewer T cell epitopes compared to the rest of this protein. When similarly illustrating the T cell recognition of SARS-CoV-2-derived peptides observed in healthy donors, we detected a comparable spread of T cell recognition in the healthy donor cohort. Interestingly, most T cell epitope clusters in the patient cohort coincide with T cell recognition in the healthy donor cohort. There are few areas that distinguish the T cell recognition observed in healthy donors from that observed in patients, these include: the C- and N-terminal regions of ORF1, parts of the N, and in general, a higher level of T cell recognition to S. In all those areas, T cell recognition in healthy donors exceed the observation from the patients (Figure 5A). Interestingly, when evaluating the prevalence of T cell responses detected towards the immunodominant epitopes identified either from the patient (Figure 1I) or the healthy donor cohort (Supplementary Figure 4C), we observed that most T cell responses dominating in patients are also detected in healthy donors, while a large fraction of epitopes domination in healthy individual are exclusively detected in this cohort (Figure 5B). This points to a substantial degree of cross-recognition to SARS-CoV-2 from pre-existing T cell populations, and that such populations might, to a large extent, drive the further expansion of T cell responses to SARS-CoV-2 infection.

**Figure 5.**
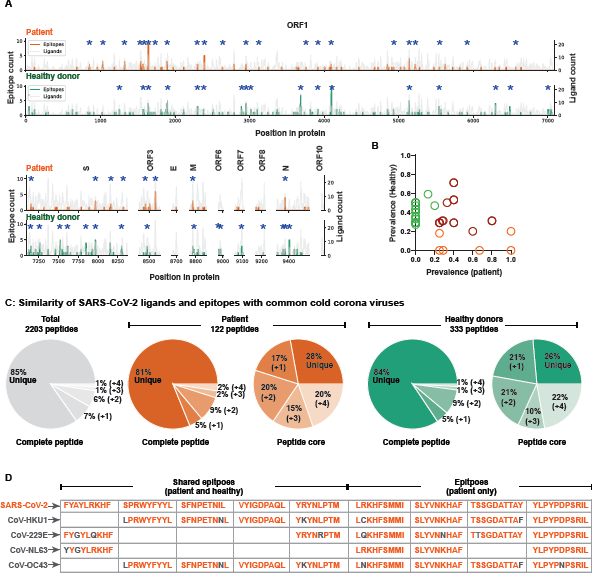
A fraction of SARS-CoV-2 epitopes shares sequence homology with widely circulating common cold coronaviruses. (**A**) SARS-CoV-2 T cell immunogenicity map across the viral proteome comparing the distribution of identified SARS-CoV-2 epitopes (patient cohort, orange line; n = 16) with the total analyzed peptides (gray line). The height of a peak indicates the number of ligands (right-Y axis) analyzed in a particular region and the number of identified epitopes (left Y-axis). The lower panel similarly maps epitopes and ligands from healthy donors (green line, n = 31). Positions significantly enriched (p-value < 0.05) with epitopes compared to the number of tested ligands are marked with asterisk. (**B**) T cell epitopes selected based on their immunodominant characteristics either in the patient (orange) or healthy donor (green) cohort, or represented in both (red) is evaluated for their T cell recognition prevalence’s in both cohorts. (**C**) Sequence similarity SARS-CoV-2 peptides with the other four common cold coronaviruses (HCoV); HCoV-HKU1, HCoV-NL63, and HCoV-229E. The gray pie chart indicates the sequence similarity of total predicted peptides from SARS-CoV-2 with any one (+1), two (+2), three (+3), or all four (+4) HCoV peptides with a variation limit of up to two amino acids within the full-length peptide. The colored pie chart shows a similar analysis for epitopes detected in the patient (n = 16) or healthy donor cohort (combined analysis of HD-1 and HD-2, N = 31) for full-length peptide and peptide core. (**D**) Examples of sequence homology for shared (between patient and healthy donors) and patient-specific T cell epitopes with one or more HCoV peptide sequence. Non-matching amino acids are shown in gray.

To further elucidate the potential origin of such a cross-reactive T cell population in the healthy donors cohort, we next evaluated the sequence homology of SARS-CoV-2 MHC-I binding peptides with the four common cold coronaviruses; HCoV-HKU1, HCoV-NL63, and HCoV-229E. With a variation limit of up to two amino acids in each peptide sequence, 15% of the total predicted peptides showed sequence similarity with one or more HCoV peptide sequence (Figure 5C, grey pie). Among the T cell recognized peptides, in both the patient and healthy donor cohorts, respectively, this fraction was comparable with 19% and 16% of T cell recognized peptides sharing sequence homology with one or more HCoV (Figure 5C). As an alternative approach the similarities were calculated by kernel method for amino acid sequences using BLOSUM62, indicating comparable sequence similarity of peptides recognized by T cells and those not recognized in reference to HCoV. However, interestingly, peptides of lowest similarity were not recognized by T cells in the patient cohort (Supplementary Figure 8).

As T cell cross-recognition can often be driven by a few key interaction points, predominantly in the ‘core’ of the peptide sequence (i.e., position 3–8) (Calis et al., 2013; Frankild et al., 2008), we restricted the sequence similarity to the core of the peptide that would most likely interact with the TCR(Bentzen et al., 2018). Based on the protein-core only, up to 74% of all the identified epitopes showed sequence homology to HCoV (one or more) (Figure 5C), suggesting these common cold viruses as a potential source of the observed low-avidity interactions in healthy donors. Further, when evaluating peptides frequently recognized by T cells in both the patients and the healthy cohort, we find evidence of substantial homology, as exemplified with the peptide sequences listed in Figure 5D. However, similar sequence homology is observed for the peptide sequences that are recognized only in the patient cohort (Figure 5D). Thus, at present, our data points to substantial T cell cross recognition being involved in shaping the T cell response to SARS-CoV-2 in COVID-19 patients, however, we find no specific enrichment of T cell recognition to peptide sequences with large sequence homology compared to the total peptide library being evaluated, and the responses that are identified in the patient samples only does not hold a more SARS-CoV-2 unique signature than those recognized in both cohorts. Interestingly, however, ORF1 being one of the elements displaying the highest T cell recognition immunogenicity, also display the highest sequence identity to HCoV (40%, as oppose to 22-34% for all other SARS-CoV-2 proteins, calculated using direct sequence alignment). Seeking to fully understand the role and origin of the underlying T cell cross-recognition will likely require an in-depth evaluation of pre- and post-infection samples.

## Discussion

We identified CD8^+^ T cell responses to 122 epitopes in 18 COVID-19 patients after screening for T cell recognition based on 3141 peptides derived from the full SARS-CoV-2 genome, and selected based on their predicted HLA-binding capacity. Of these, a few dominant T cell epitopes were recognized in the majority of the patients. Strikingly, both dominant and subdominant T cell epitopes were cross-recognized by low-level existing T cell populations in SARS-CoV-2 unexposed healthy individuals. We have observed that the SARS-CoV-2 dominant epitopes mount very strong T cell responses, with up to 25% of all CD8^+^ lymphocytes specific to a single epitope (two overlapping peptides with same peptide core).

Pre-existing immunity based on cross-reactive T cells can influence how our immune system reacts upon viral exposure. One way could be via faster expansion of pre-existing memory cells upon initial exposure to viral infection. A similar outcome and benefit of pre-existing T cell immunity have been shown in the case of flu pandemic virus H1N1 (Sridhar et al., 2013; Wilkinson et al., 2012). Additionally, hyperactivation of pre-existing T cells can contribute to short- and long-term disease severity via inflammation and autoimmunity, as increased production of IFN-γ by CD4^+^ and CD8^+^ T cells has been observed in severe COVID-19 patients (Wang et al., 2020). Furthermore, it has been reported (Ehrenfeld et al., 2020) that SARS-CoV-2 infection can be a triggering factor for autoimmune reactions and severe pneumonia with sepsis leading to acute respiratory distress syndrome (ARDS), bone-marrow affection with pancytopenia and organ-specific autoimmunity (Gagiannis et al., 2020; Henderson et al., 2020; Hersby et al., 2020). Importantly, pre-existing T cell immunity can influence vaccination outcomes, as they can induce a faster and better immune response. The ORF1 protein regions are highly conserved within coronaviruses (Cui et al., 2019), and show the highest HCoV identity among SARS-CoV-2 proteins; and most of the immunodominant epitopes that we have identified belong to the ORF1 region. Thus, a detailed evaluation of these T cell epitopes could be of value in vaccine design.

Most vaccine-development efforts are currently focusing on mounting antibody-responses to the spike protein, with limited focus on T cell immunity. However, studies point to a potential fast decline in antibody titers after infection (Seow et al., 2020; Vabret, 2020), and such kinetics might be similar following vaccinations. Thus, the involvement of T cell immunity might be a relevant focus if antibody titers cannot sufficiently protect against infections. In such scenarios, T cell immunity can sustain the antibody responses and provide a direct source of T cells clearing virus-infected cells. For the involvement of T cell immunity in vaccine development, our data suggest that the inclusion of other virus proteins, such as ORF1 or 3, might be highly relevant.

T cell recognition of SARS-CoV-2-derived peptides in both COVID-19 patients and healthy donors has prompted us to understand the role of T cell cross-reactivity in controlling infections. In recent years, technology improvements in TCR characterization have allowed us to interrogate the TCR-pMHC interaction from a structural approach, while obtaining experimental information related to the peptide amino acids that are crucial to T cell recognition (Adams et al., 2016; Bentzen and Hadrup, 2019; Birnbaum et al., 2014; Garcia et al., 1996, 1998; Linette et al., 2013). Such efforts have taught us that T cell cross-recognition is very difficult to predict, without knowing the precise interaction required for the given TCR, as even T cell epitopes with as low as 40% sequence homology can be recognized by a given TCR (Bentzen et al., 2018). Therefore, the underlining source of T cell cross-reactivity might arise from a larger set of epitopes within the HCoV viruses, including sequences with larger variation than those evaluated here (i.e., max. two amino acid variants per peptide sequence/peptide core).

While the T cell recognition itself was largely overlapping in identity between patients and healthy donors, the magnitude for the T cell responses and, in particular, the T cell phenotype of SARS-CoV-2-specific T cells was substantially different. A unique phenotype characterization demonstrated a strong activation profile of SARS-CoV-2-specific T cells only in patients. This strong ‘activation signature’ (high expression of CD38, CD39, CD69, PD1) was further enhanced in patients requiring hospitalization. Such strong and highly activated T cell responses should be able to clear the virus, and hence our data further support the notion that some severely affected patients might suffer from hyperactivation of their T cell compartment as a consequence of their primary viral infection, which may even be cleared.

Taken together, the data presented here demonstrate a substantial role for T cell recognition in COVID-19, and in-depth evaluations in larger cohorts over time will provide essential insight to the role of such T cells in disease severity and how pre-existing T cell immunity can be leveraged to fight novel infections.

## Methods

### Clinical samples

Approval for the study design and sample collection was obtained from the Commitee on Health Research Ethics in the Capital Region of Denmark. All included patients and health care employees gave their informed written consent for inclusion. PBMC samples from 18 COVID-19 infected patients were used in this study. Blood samples were collected as close as possible to the first COVID-19 positive test. The patient cohort included samples from individuals with severe symptoms who required hospital care (hospitalized; n = 11) and patients with mild symptoms not requiring hospital care (outpatient; n = 7). For hospitalized patients, we collected full data from the medical record regarding disease course, age, gender, travel history, performance status, symptoms, comorbidity, medications, laboratory findings, diagnostic imaging, treatment, need of oxygen, need for intensive care, and an overall estimate of disease severity (Supplementary Table 2). For outpatients, we used a questionnaire to collect data on comorbidity, travel history, medications, and performance status.

COVID-19 infection was diagnosed by one of the four platforms; BGI (BGI Covid-19 RT-PCR kit), PantherFusion (Hologic), Roche Flow (Roche MagNA Pure 96, Roche LightCycler 480 II real-time PCR), and Qiaflow (QIAsymphony or RotorGene, Qiagen). In the last three platforms, LightMix Modular SARS-CoV (COVID-19) E-gene (# 53-0776-96) has been used. The diversity of platforms used were due to supply issues. All platforms were validated using validation kits and panels from the Statens Serum Institute (SSI), Denmark. Most patients had more than one positive test for COVID-19. Swabs, sputum, and tracheal secretion were used depending on the setting.

For the pre-COVID-19 healthy donor cohort (n = 18), we used samples collected prior to October 2019 and obtained from the central blood bank, Rigshospitalet, Copenhagen, in an anonymized form. Additionally, we included 20 health care employees from Herlev Hospital during the COVID-19 pandemic, who were at high risk of COVID-19 infection but not positive, as a cohort to follow immune responses in a potentially exposed population.

PBMCs from all three cohorts were isolated immediately after sampling using Ficoll-Paque PLUS (GE Healthcare) density gradient centrifugation and were cryopreserved thereafter at a density of 2–20 × 10^6^ cells/mL.

### SARS-CoV-2 peptide selection

Potential HLA class I binding peptides were predicted from the complete set of 8–11mer peptides contained within the Wuhan seafood market pneumonia virus isolate Wuhan-Hu-1 (GenBank ID MN908947.3) to a set of ten prevalent and functionally diverse HLA molecules (HLA-A01:01, HLA-A02:01, HLA-A03:01, HLA-A24:02, HLA-B07:02, HLA-B08:01, HLA-B15:01 HLA-C06:02, HLA-C07:01, HLA-C07:02) using a preliminary version of NetMHCpan-4.1 (http://www.cbs.dtu.dk/services/NetMHCpan/index_v0.php_v0.php)[PMID: 32406916]. For peptides predicted from ORF1 protein, a percentile rank binding threshold of 0.5% was used, and for peptides derived from all other proteins, a threshold of 1% was used. Altogether, 2203 peptides were selected, binding to one or more HLA molecules, generating 3141 peptide-HLA pairs for experimental evaluation (Supplementary Table 1). All peptides were custom synthesized by Pepscan Presto BV, Lelystad, The Netherlands. Peptide synthesis was done at a 2 μmol scale with UV and mass spec quality control analysis for 5% random peptides by the supplier.

### MHC class I monomer production

All ten MHC-I monomer types were produced using methods previously described (Saini et al., 2013). Briefly, MHC-I heavy chain and human ß2-microglobulin (hß2m) were expressed in *Escherichia coli* using pET series expression plasmids. Soluble denatured proteins of the heavy chain and hß2m were harvested using inclusion body preparation. The folding of these molecules was initiated in the presence of UV labile HLA specific peptide ligands (Hadrup et al., 2009a). HLA-A02:01 and A24:02 molecules were folded and purified empty, as described previously (Saini et al., 2019). Folded MHC-I molecules were biotinylated using the BirA biotin-protein ligase standard reaction kit (Avidity, LLC-Aurora, Colorado), and MHC-I monomers were purified using size exclusion chromatography (HPLC, Waters Corporation, USA). All MHC-I folded monomers were quality controlled for their concentration, UV degradation, and biotinylation efficiency, and stored at −80°C until further use.

### DNA-barcoded multimer library preparation

The DNA-barcoded multimer library was prepared using the method developed by Bentzen et al. (Bentzen et al., 2016). Unique barcodes were generated by combining different A and B oligos, with each barcode representing a 5’ biotinylated unique DNA sequence. These barcodes were attached to phycoerythrin (PE) or allophycocyanin (APC) and streptavidin-conjugated dextran (Fina BioSolutions, Rockville, MD, USA) by incubating them at 4°C for 30 min to generate a DNA-barcode-dextran library of 1325 unique barcode specificities. SARS-CoV-2 pMHC libraries were generated by incubating 200 μM peptide of each peptide with 100 μg/mL of respective MHC molecules for 1 h using UV-mediated peptide exchange (HLA-A01:01, A03:01, B07:02, B08:01, B15:01, C06:02, C07:01, and C07:02) or direct binding to empty MHC molecules (HLA-A02:01 and A24:02). HLA-specific DNA-barcoded multimer libraries were then generated by incubating pMHC monomers to their corresponding DNA barcode dextrans at 4°C for 30 min, thus providing a DNA barcode-labeled dextran for each peptide-MHC (pMHC multimer) specifically to probe respective T cell population. A similar process was followed to generate DNA-barcoded pMHC multimers for CEF epitopes using APC- and streptavidin-conjugated dextran attached with unique barcodes.

### T cell staining with DNA-barcoded pMHC multimers and phenotype panel

All COVID-19 patient and healthy donor samples were HLA genotyped for HLA-A, B, and C loci (IMGM Laboratories GmbH, Germany, next-generation sequencing) (Supplementary Table 7). Patient and healthy donor HLA-matching SARS-CoV2 and CEF pMHC multimer libraries were pooled (as described previously (Bentzen et al., 2016)) and incubated with 5–10 × 10^6^ PBMCs (thawed and washed twice in RPMI + 10% FCS, and washed once in barcode cytometry buffer) for 15 min at 37°C at a final volume of 60 μL. Cells were then mixed with 40 μL of phenotype panel containing surface marker antibodies (Supplementary Table 3) and a dead cell marker (LIVE/DEAD Fixable Near-IR; Invitrogen L10119) (final dilution 1/1000), and incubated at 4°C for 30 min. Cells were washed twice with barcode cytometry buffer and fixed in 1% PFA.

Cells fixed after staining with pMHC-multimers were acquired on a FACSAria flow cytometer instrument (AriaFusion, Becton Dickinson) and gated by the FACSDiva acquisition program (Becton Dickinson), and all the PE-positive (SARS-CoV-2 multimer binding) and APC-positive (CEF multimer binding) cells of CD8^+^ gate were sorted into pre-saturated tubes (2% BSA, 100 μl barcode cytometry buffer) (Supplementary Figure 9A). Sorted cells belonging to each sample were then subjected to PCR amplification of its associated DNA barcode(s). Cells were centrifuged for 10 min at 5000 × g, and the supernatant was discarded with minimal residual volume. The remaining pellet was used as the PCR template for each of the sorted samples and amplified using the Taq PCR Master Mix Kit (Qiagen, 201443) and sample-specific forward primer (serving as sample identifier) A-key36. PCR-amplified DNA barcodes were purified using the QIAquick PCR Purification kit (Qiagen, 28104) and sequenced at PrimBio (USA) using the Ion Torrent PGM 314 or 316 chip (Life Technologies).

### DNA-barcode sequence analysis and identification of pMHC specificities

To process the sequencing data and automatically identify the barcode sequences, we designed a specific software package, ‘Barracoda’ (https://services.healthtech.dtu.dk/service.php?Barracoda-1.8). This software tool identifies the barcodes used in a given experiment, assigns sample ID and pMHC specificity to each barcode, and calculates the total number of reads and clonally reduced reads for each pMHC-associated DNA barcode. Furthermore, it includes statistical processing of the data. Details are given in Bentzen et al. (Bentzen et al., 2016). The analysis of barcode enrichment was based on methods designed for the analysis of RNA-seq data and was implemented in the R package edgeR. Fold changes in read counts mapped to a given sample relative to mean read counts mapped to triplicate baseline samples were estimated using normalization factors determined by the trimmed mean of M-values. P-values were calculated by comparing each experiment individually to the mean baseline sample reads using a negative binomial distribution with a fixed dispersion parameter set to 0.1 (Bentzen et al., 2016). False-discovery rates (FDRs) were estimated using the Benjamini-Hochberg method. Specific barcodes with an FDR < 0.1% were defined as significant, determining T cell recognition in the given sample. At least 1/1000 reads associated with a given DNA barcode relative to the total number of DNA barcode reads in that given sample was set as the threshold to avoid false-positive detection of T cell populations due to the low number of reads in the baseline samples. T cell frequency associated with each significantly enriched barcode was measured based on the % read count of the associated barcode out of the total % multimer-positive CD8^+^ T cells population. In order to exclude potential pMHC elements binding to T cells in a non-specific fashion, non-HLA-matching healthy donor material was included as a negative control. Any T cell recognition determined in this samples was subtracted from the full data set.

### T cell expansion and combinatorial tetramer staining

PBMCs from healthy donors were expanded *in-vitro* using pMHC-dextran complexes conjugated with SARS-CoV-2-derived peptides and cytokines (IL-2 and IL-21) for 2 weeks either with single pMHC specificity or with a pool of up to ten pMHC specificities. PBMCs were expanded for 2 weeks in X-vivo media (Lonza, BE02-060Q) supplemented with 5% human serum (Gibco, 1027-106). Expanded cells were used to measure peptide-specific T cell activation or stained using pMHC tetramers to detect T cells recognizing SARS-CoV-2 epitopes.

*In-vitro* expanded healthy donor PBMCs were examined for SARS-CoV-2 reactive T cells using combinatorial tetramer staining (Sick Andersen et al., 2012). Individual HLA-restricted pMHC complexes were generated using direct peptide loading (HLA-A02:01 and A24:02) or UV-mediated peptide exchange (all other HLAs) as described above and conjugated with fluorophore-labeled streptavidin molecules. For 100 μL pMHC monomers, 9.02 μL (0.2 mg/mL stock, SA-PE-CF594 (Streptavidin - Phycoerythrin/CF594; BD Biosciences 562318), SA-APC (Biolegend 405207)) or 18.04 μL (0.1 mg/mL stock, SA-BUV395 (Brilliant Ultraviolet 395; BD 564176), SA-BV421 (Brilliant Violet 421; BD 563259), and SA-BV605 (Brilliant Violet 605; BD 563260)) of streptavidin conjugates were added and incubated for 30 min at 4°C, followed by addition of D-biotin (Sigma) at 25 μM final concentration to block any free binding site. pMHC tetramers for each specificity were generated in two colors by incubating pMHC monomers and mixed in a 1:1 ratio before staining the cells. Expanded cells were stained with 1 μL of pooled pMHC multimers per specificity (in combinatorial encoding) by incubating 1–5 × 10^6^ cells for 15 min at 37°C in 80 μL total volume. Cells were then mixed with 20 μL antibody staining solution CD8-BV480 (BD B566121) (final dilution 1/50), dump channel antibodies (CD4-FITC (BD 345768) (final dilution 1/80), CD14-FITC (BD 345784) (final dilution 1/32), CD19-FITC (BD 345776) (final dilution 1/16), CD40-FITC (Serotech MCA1590F) (final dilution 1/40), CD16-FITC (BD 335035) (final dilution 1/64)) and a dead cell marker (LIVE/DEAD Fixable Near-IR; Invitrogen L10119) (final dilution 1/1000) and incubated for 30 min at 4°C. Cells were then washed twice in FACS buffer (PBS+2% FCS) and acquired on a flow cytometer (Fortessa, Becton Dickinson). Data were analyzed using FlowJo analysis software.

### T cell functional analysis

For functional evaluation of T cells from PBMCs of COVID-19 patients or PBMCs expanded from healthy donors, 1–2 × 10^6^ cells were incubated with 1 μM of SARS-CoV-2 minimal epitope or pool of up to ten epitopes (1 μM/peptide) for 9 h at 37°C in the presence of protein transport inhibitor (GolgiPlug; BD Biosciences, 555029; final dilution 1/1000). Functional activation of T cells was measured using intracellular cytokines IFN-γ (BD Bioscience, 341117; final dilution 1/20) and TNF-α (Biolegend, 502930; final dilution 1/20). Cells incubated with Leukocyte Activation Cocktail (BD Biosciences, 550583; final dilution 1/500) were used as a positive control, and HLA-specific irrelevant peptides were used as negative controls. Surface marker antibodies CD3-FITC (BD Biosciences 345764 (final dilution 1/20)), CD4-BUV395 (BD Biosciences 742738 (final dilution 1/300), CD8-BV480 (BD Biosciences B566121 (final dilution 1/50)), and dead cell marker (LIVE/DEAD Fixable Near-IR; Invitrogen L10119) (final dilution 1/1000)) were used to identify CD8^+^ T cells producing intracellular cytokines (Gating strategy, Supplementary Figure 5).

### Flow cytometry analysis and UMAP

For phenotype analysis, all samples were analyzed using FlowJo data analysis software (FlowJO LLC). Frequencies of specific cell populations were calculated according to the gating strategy shown in Supplementary Figure 9B. For combinatorial tetramers staining, T cell binding to specific-pMHC tetramers was identified using the gating plan described in the original study (Hadrup et al., 2009b). For UMAP analysis (McInnes et al., 2018), FCS files of samples from the patient and healthy cohorts were concatenated (160,000 total cells), downsampled (FlowJo plugin), and visualized using UMAP (Version 2.2, FLowJo plugin) analysis based on the selected markers; CD3, CD4, CD8, CD38, CD39, CD69, CD137, HLA-DR, PD-1, CCR7, CD45RA, CD27, and CD57.

### Sequence homology analyses

To evaluate the homology between SARS-Cov2 and HCoV, both epitopes (peptides recognized by T cells) and ligands (peptide not recognized by T cells) were mapped to their respective source protein from the SARS-CoV-2 proteome. Enrichment analysis of the epitopes in the given region of the proteins was based on testing whether the count of observed epitopes exceeded what we expected from the number of ligands tested at each position. Epitopes were considered successes and the count of ligands were regarded as the number of trials in a binomial test. The probability of success was derived from the average ratio of epitope to ligand per position across each protein. The test was ‘one-sided’ with a significance level at 0.05.

The similarity of SARS-CoV-2 ligands and epitopes from both patient and healthy donor cohorts to a set of human common cold corona viruses (HCoV-HKU1, HCoV-229E, HCoV-NL63, HCoV-OC43) was tested using two methods. The first approach utilized a kernel method for amino acid sequences using BLOSUM62 (Shen et al., 2012). The second approach was a simple string search allowing up to two mismatches. Based on the second approach each epitope was categorized by how many, if any, of the common cold viruses it would match with. Both methods were applied to the full peptide length and to the peptide core.

### Data processing and statistics

T cell recognition was determined based on the DNA-barcoded pMHC multimer analysis and evaluated through ‘baracoda’ (see above). The data was plotted using python 3.7.4. For all plots, peptide sequences with no significant enrichments are shown as gray dots and all peptide with a negative enrichment are set to LogFc equal zero (Figure 1D, E, G; Figure 2A, B; Supplementary Figure 2). Box plots for data quantification and visualization were generated, and their related statistical analyses were performed using GraphPad Prism (GraphPad Software Inc.) (Figure 2C; Figure 3C, D, F; Figure 4A, B, D) or R studio (Supplementary Figure 8). For unpaired comparisons Mann-Whitney test was done using GraphPad Prism, all p values are indicated in figure legends. Flow Cytometry data were analyzed using FlowJo (version 10). Immunogenicity scores (Figure 1F, H; Supplementary Figure 4) were calculated (as %) by dividing total identified T cell reactivity associated with a HLA or protein with the total number of specificities analyzed in a given cohort (number of peptides multiplied by number of patient with a given HLA). Staining index (Figure 2C) was calculated as; ((mean fluorescence intensity (MFI) of multimer^+^ cells – MFI of multimer^-^ cells)/2 × standard deviation (SD) of multimer^-^ cells)). MFI of multimer^+^ and multimer^-^ CD8^+^ T cells and the SD of the multimer^-^ CD8^+^ T cells from FlowJo analysis for patient and healthy donor samples.

### Data and code availability

The data that support the finding of this study, in addition to the supplementary supporting data, and the code used to generate the plots and analyses can be accessed from the corresponding author upon reasonable request.

## Supplementary data

1. **Supplementary Figure 1. Details of SARS-CoV-2-derived peptides in relation to their HLA restriction, and coverage in the COVID-19 patients analyzed in this study.**
2. **Supplementary Figure 2. Example of genome-wide screening for SARS-CoV-2-reactive T cells in individual patient samples.**
3. **Supplementary Figure 3. Functional validation of SARS-CoV-2-specific T cell responses identified in COVID-19 patient samples.**
4. **Supplementary Figure 4. Summary of SARS-CoV-2-specific T cell reactivity identified in healthy donors.**
5. **Supplementary Figure 5. Gating strategy and functional evaluation of SARS-CoV-2 reactive CD8^+^ T cells identified in healthy donors.**
6. **Supplementary Figure 6. Summary of CEF-specific T cell responses identified in patient and healthy donors and their phenotype comparison.**
7. **Supplementary Figure 7. Memory phenotype of SARS-CoV-2 reactive CD8^+^ T cells.**
8. **Supplementary Figure 8. Sequence similarities of SARS-CoV-2 peptides with HCoV.**
9. **Supplementary Figure 9. Gating strategy for sorting multimer^+^ CD8^+^ T cells and phenotype analysis.**
10. **Supplementary Table 1: List of SARS-CoV-2 peptides with their HLA rank score.**
11. **Supplementary Table 2: COVID-19 patient details.**
12. **Supplementary Table 3: Phenotype antibody panel.**
13. **Supplementary Table 4: CEF epitope library.**
14. **Supplementary Table 5: SARS-CoV-2-specific T cell epitopes identified in COVID-19 patients.**
15. **Supplementary Table 6a: SARS-CoV-2-specific T cell epitopes identified in pre-COVID-19 healthy donors (HD-1)**
16. **Supplementary Table 6b: SARS-CoV-2-specific T cell epitopes identified in high-risk healthy donors (HD-2)**
17. **Supplementary Table 7: HLA genotype data for patient and healthy donors**

## Acknowledgements

We thank all patients and healthy donors for participating and contributing the analyzed samples; Bente Rotbøl, Anna Gyllenberg Burkal, Anni Flarup Løye, and Pia Taaning Petersen for their excellent technical support; The clinical research unit for patient inclusion – Kathrine Friser Kokholm, Mette Lundø Sieg, Jytte Kock; Department of microbiology Herlev Hospital Lene Nielsen, and all the collaborators for their active participation to this work.

## Funding

The work is supported by the Independent Research Fund Denmark (grant no. 0214-00053B, 2020).

## Author contributions

S.K.S. conceived the idea, designed and performed experiments, analyzed data, prepared figures, and wrote the manuscript. D.S.H. designed research, facilitated patient samples, and wrote the manuscript. T.T designed research, performed experiments, and analyzed data. H.R.P. analyzed data, performed bioinformatics analysis, prepared figures, and wrote the manuscript. S.P.A.H. performed the experiments. M.N. supervised and performed the research, and wrote the manuscript. A.O.G. conceived the idea, supervised clinical study; patient participation, data, and sample collection, and wrote the manuscript, S.R.H. conceived the idea, designed research, wrote the manuscript, and supervised research.

## Competing interests

S.R.H. and S.K.S. are cofounders of Teramer Shop, S.R.H. is a cofounder of PokeAcell. Commercialization of DNA-barcoded technology is licensed to Immudex. These activities pose no competing interests related to the data reported here. All other authors declare no competing interests.

