## Supplementary material for "SARS-CoV-2 genome-wide mapping of CD8 T cell recognition reveals strong immunodominance and substantial CD8 T cell activation in COVID-19 patients"

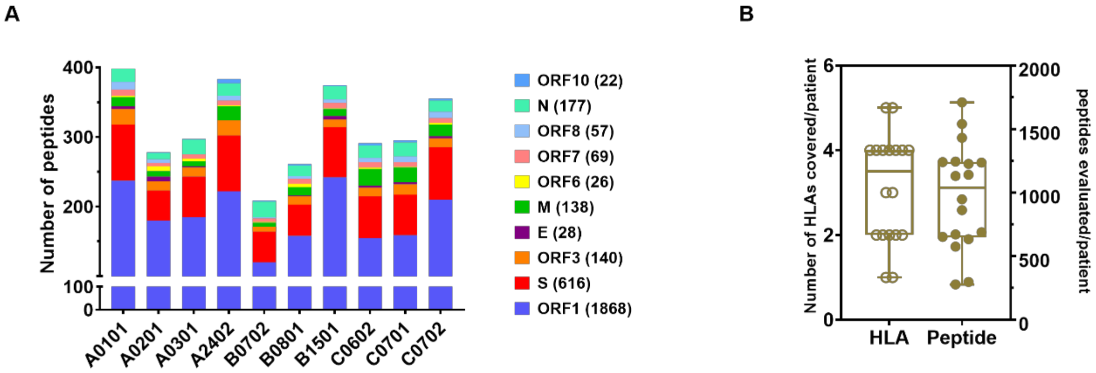


**Supplementary Figure 1. Details of SARS-CoV-2-derived peptides in relation to their HLA restriction, and coverage in the COVID-19 patients analyzed in this study. (A)** Bar plots showing the distribution of SARS-CoV-2 peptides and their HLA-restriction (3141 peptide-HLA pairs) across the viral genome. Binding prediction of the 8–11 amino acid peptides for the ten listed HLAs was made using NetMHCpan4.1. Total peptide-MHC specificities analyzed for each protein are shown in parentheses next to the respective SARS-CoV-2 protein. **(B)** Box plots showing the number of peptide-MHC specificities (right Y-axis) evaluated for $CD8^+$ T cell reactivity in COVID-19 patients based on the HLA expression of individual patients matching the ten selected HLAs, i.e. number of HLAs covered/patient (left Y-axis). Numbers; HLA coverage = 3.1 ± 1.2, peptides evaluated = 972 ± 414 (n = 18, mean ± SD).

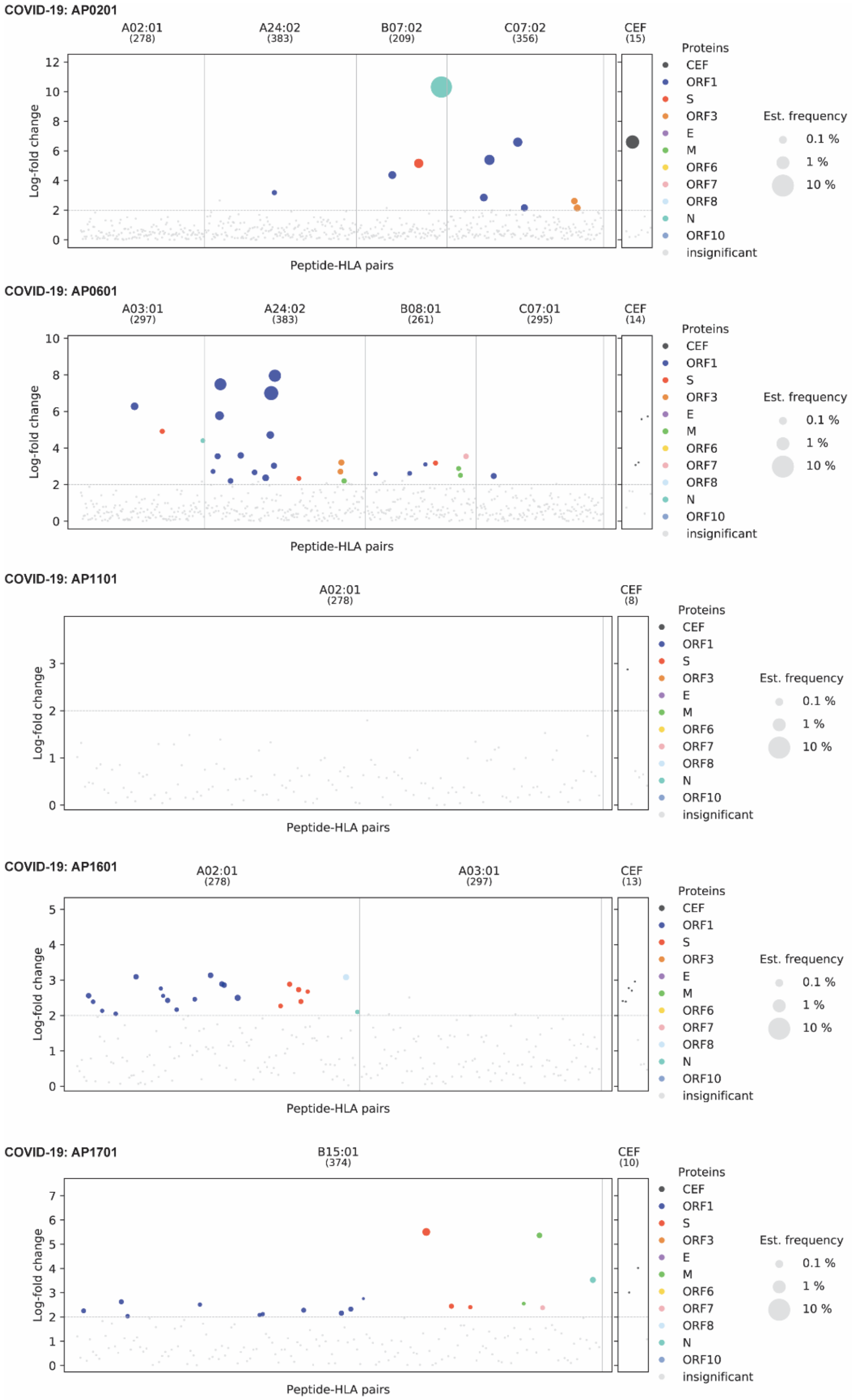


**Supplementary Figure 2. Example of genome-wide screening for SARS-CoV-2-reactive T cells in individual patient samples.** CD8$^+$ T cell recognition to individual epitopes were identified based on the enrichment of DNA barcodes associated with each of the tested peptide specificities (LogFc>2 and p < 0.001, *barracoda*). Significant T cell recognition of individual peptide sequences are colored based on their protein of origin and segregated based on their HLA-specificity. The size of the dot represents the estimated frequency of pMHC multimer positive CD8$^+$ T cell populations for each of the recognized epitopes. The black dots show CD8$^+$ T cells reactive to CEF peptides. All peptide sequences with no significant enrichments are shown as gray dots. Representative examples from 5 different COVID19 patients are depicted.

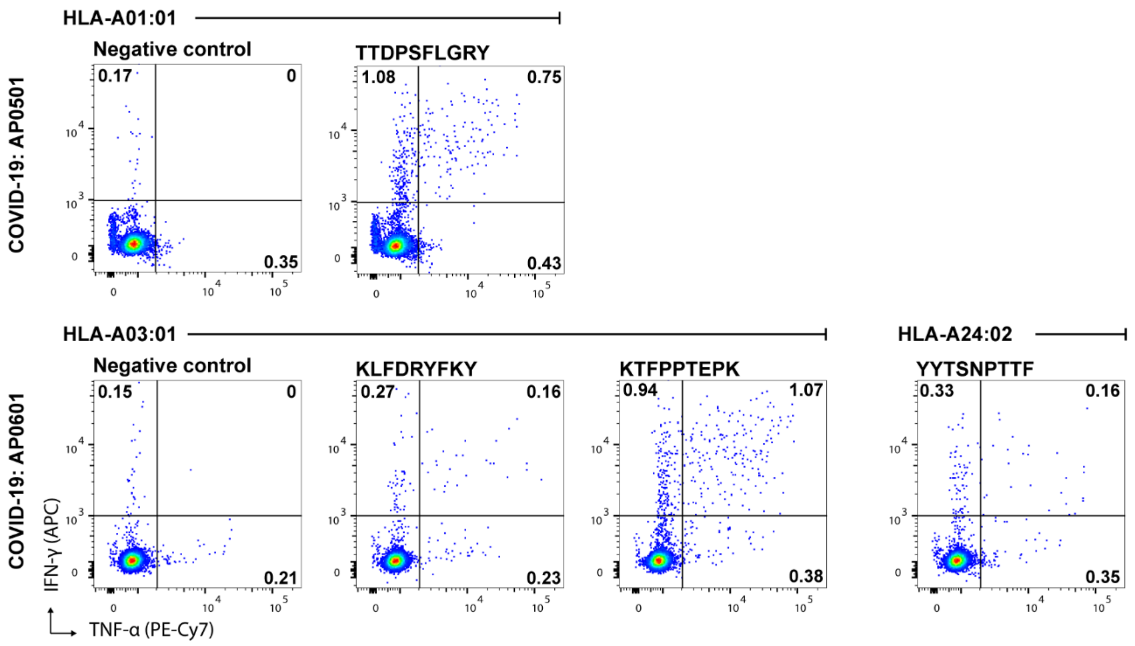


**Supplementary Figure 3. Functional validation of SARS-CoV-2-specific T cell responses identified in COVID-19 patient samples.** Flow cytometry plots of intracellular cytokine staining of PBMCs pulsed with selected SARS-CoV-2-derived peptides that were identified as T cell epitopes in the COVID-19 patients using the DNA-barcoded multimer analysis (**Supplementary Table 5**). PBMCs from two COVID-19 patients were analyzed for functional activation after stimulation with indicated peptides or with a HLA-matching irrelevant peptide (negative control). The numbers on the plot indicate the frequency (%) of $CD8^+$ T cells positive for the analyzed cytokines, IFN-$\gamma$ and TNF-α. The gating strategy of the flow cytometry analysis is shown in **Supplementary Figure 5A**.

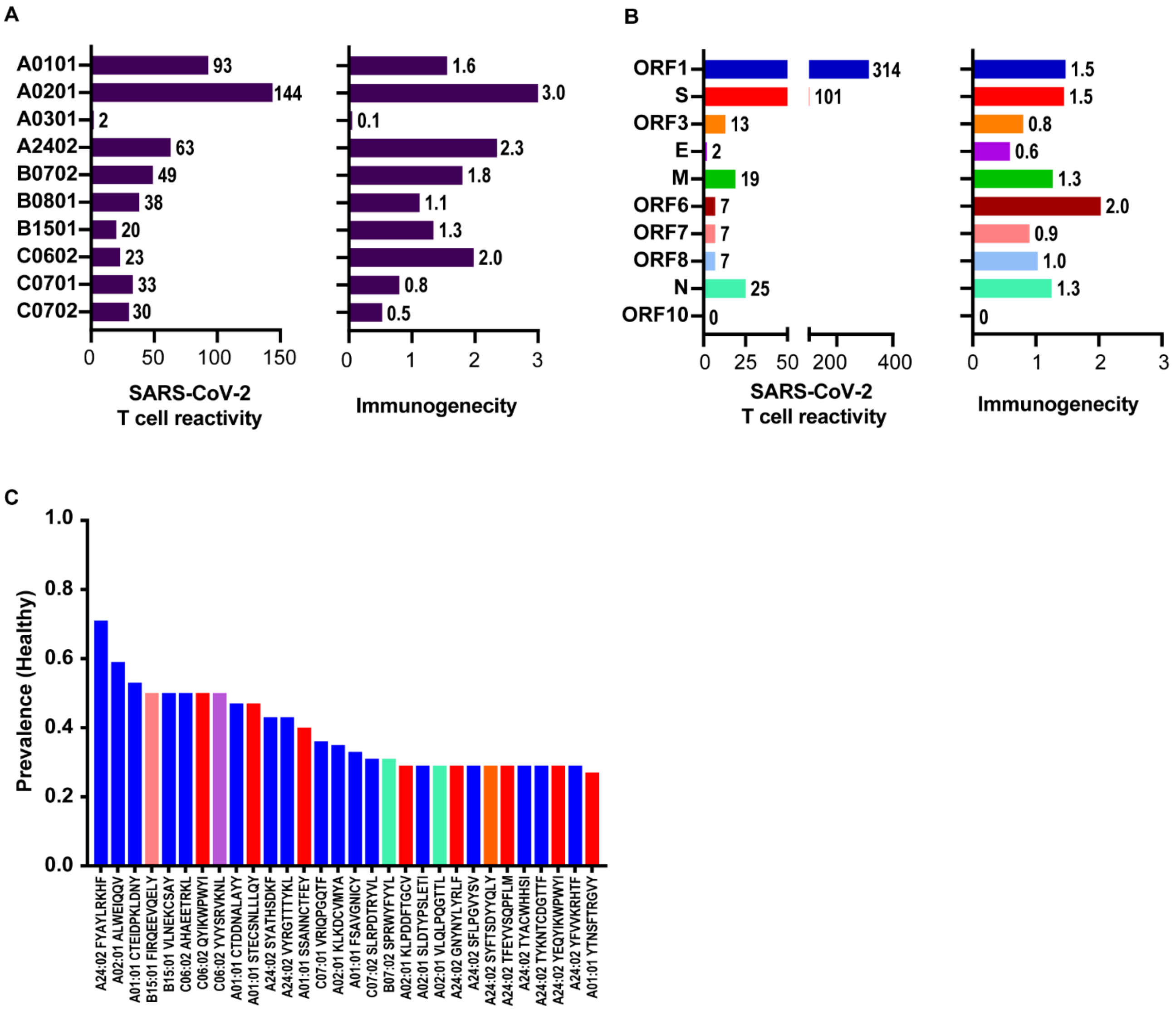


**Supplementary Figure 4. Summary of SARS-CoV-2-specific T cell reactivity identified in healthy donors. (A)** HLA-specific T cell recognition of SARS-CoV-2-derived peptides and HLA-restricted immunogenicity score (%) in healthy donors (pooled data of HD-1 and HD-2 cohorts, plots shown in Figure 2A, B). (**B**) Similar to A, T cell recognition shown based on the protein of origin and their immunogenicity score (%). (**C**) Bar plots showing prevalence (fraction of analyzed samples) of immunodominant epitopes (detected in two or more donors and >25% of the tested samples for a given HLA molecules) in healthy donors.

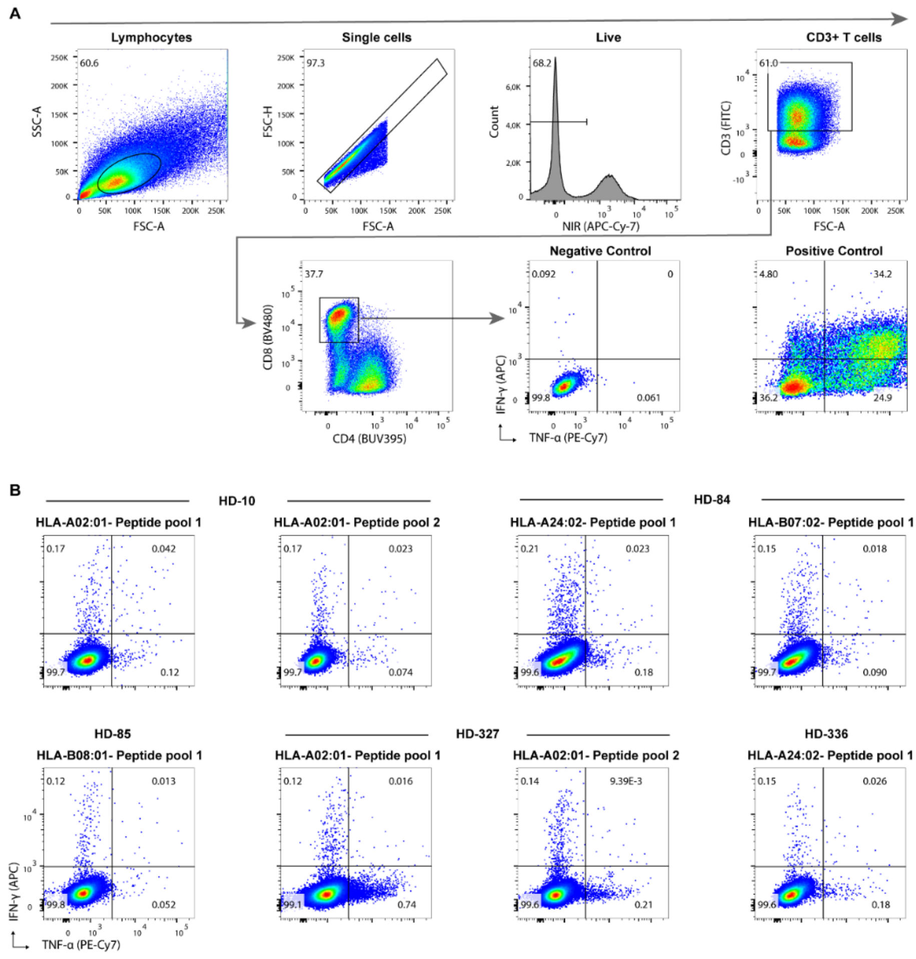


**Supplementary Figure 5. Gating strategy and functional evaluation of SARS-CoV-2 reactive CD8$^{+}$ T cells identified in healthy donors. (A)** Representative flow cytometry plots for gating strategy used to measure cytokines (IFN-γ, and TNF-α) in CD8$^{+}$ T cells as a measure of functional activation upon stimulation with specific peptides. The numbers on the plots show the frequency of the gated cells (% of the parent population). Cells incubated with an irrelevant peptide or no peptide were used as a negative control, and cells incubated with a leukocyte activation cocktail (containing Phorbol 12-Myristate 13-Acetate, Ionomycin, and the protein transport inhibitor Brefeldin A) were used as a positive control. **(B)** Flow cytometry plots of PBMCs from several healthy donors (Pre-COVID-19, from HD-1 cohort) expanded for 2 weeks based on pMHC specific stimulation and evaluated for functional T cell activation upon stimulation with pool SARS-CoV-2-derived peptides. Expanded cells were incubated with a pool of up to ten peptides restricted to a given HLA depending on the SARS-CoV-2 T cell reactivates identified in the DNA-barcoded multimer analysis of the healthy donors (**Supplementary Table 5**). The relevant negative control is depicted in (A).

**A**

**COVID-19 patients**

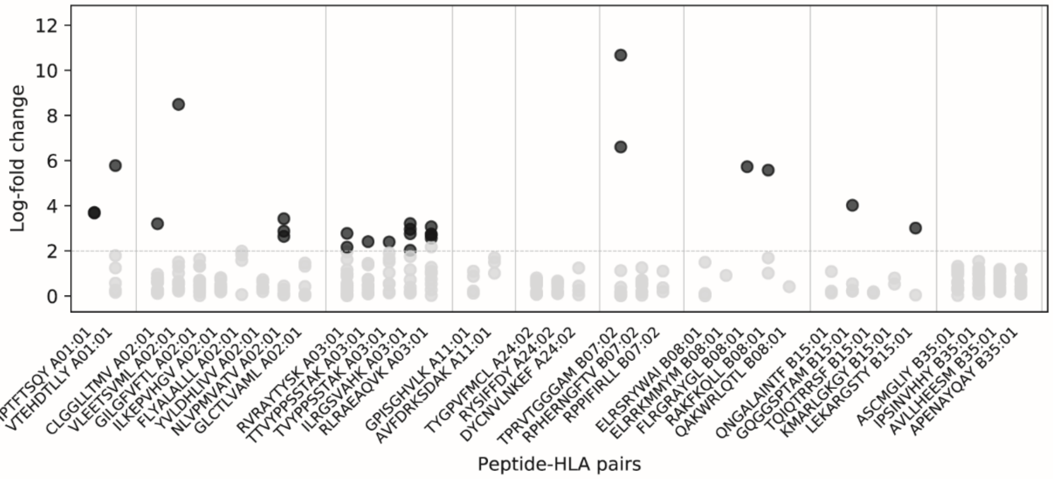


**Healthy donors: Pre-COVID-19 (HD-1)**

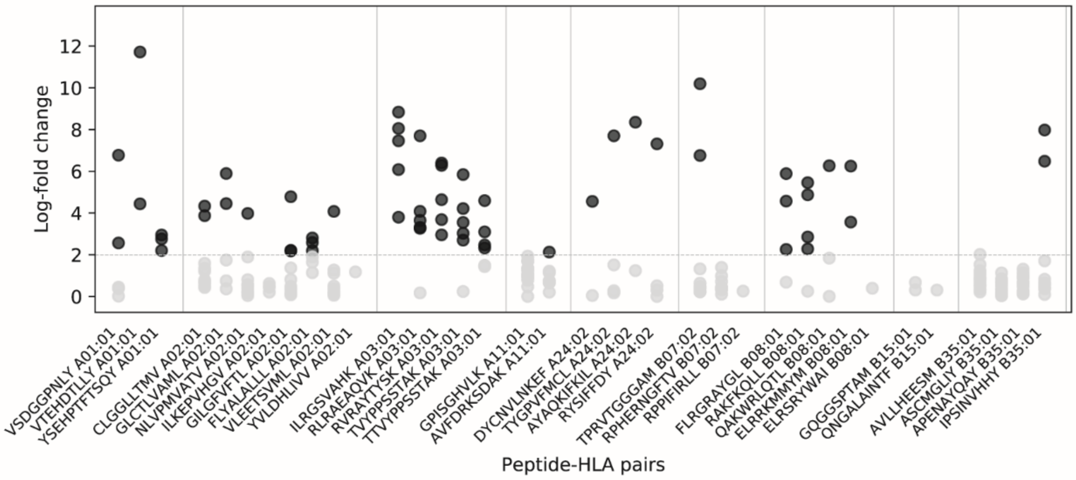


**Healthy donor: Hospital staff (HD-2)**

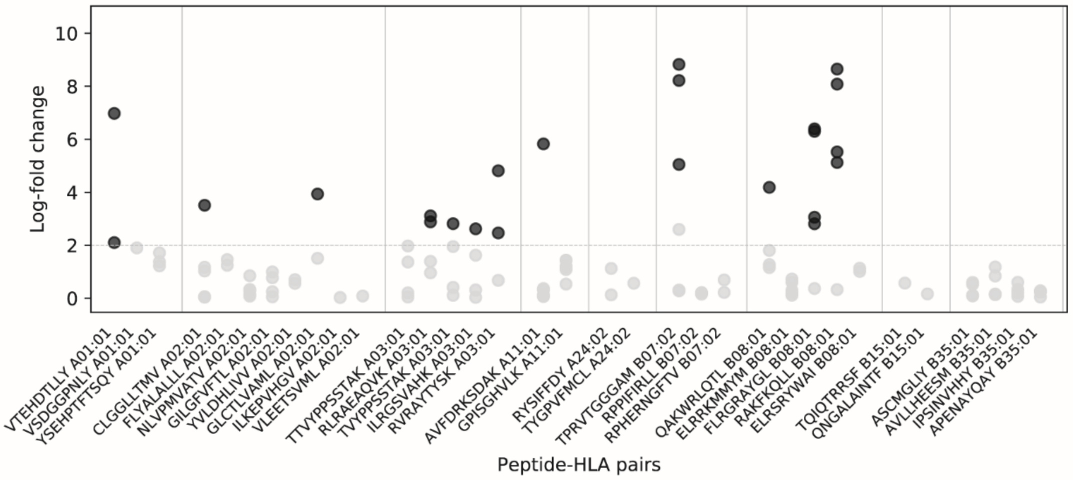


**B**

**CEF multimers**

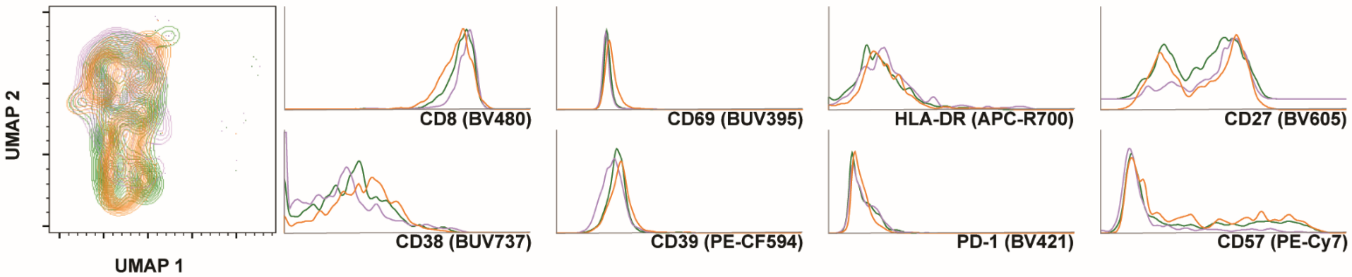

**Supplementary Figure 6. Summary of CEF-specific T cell responses identified in patient and healthy donors and their phenotype comparison. (A)** Each plot summarize CEF-specific $CD8^+$ T cell identified in the patient cohort and the two healthy donor cohorts. Together with SARS-CoV-2-derived peptides, DNA-barcoded pMHC-multimers of 39 CEF (CMV, EBV, and Influenza) epitopes were included in the analysis (**Supplementary Table 4**). $CD8^+$ T cell recognition to individual epitopes were identified based on the enrichment of DNA barcodes associated with each of the tested peptide specificities (LogFc>2 and $p < 0.001$, *barracoda*). Significant T cell recognition of individual peptide sequences are shown in black circles. Multiple circles of the same pMHC specificity indicate T cell responses identified in different samples of the cohort. No significant enrichments are shown as gray dots. The X-axis is labeled with peptide sequence and HLA restriction for a given CEF epitope. **(B)** UMAP overlay of CEF $multimer^+$ (APC) $CD8^+$ T cells identified in patient and healthy donors. Histograms compare phenotype markers expressed by CEF-specific CD8 T cells in patients and the two healthy donor cohorts.

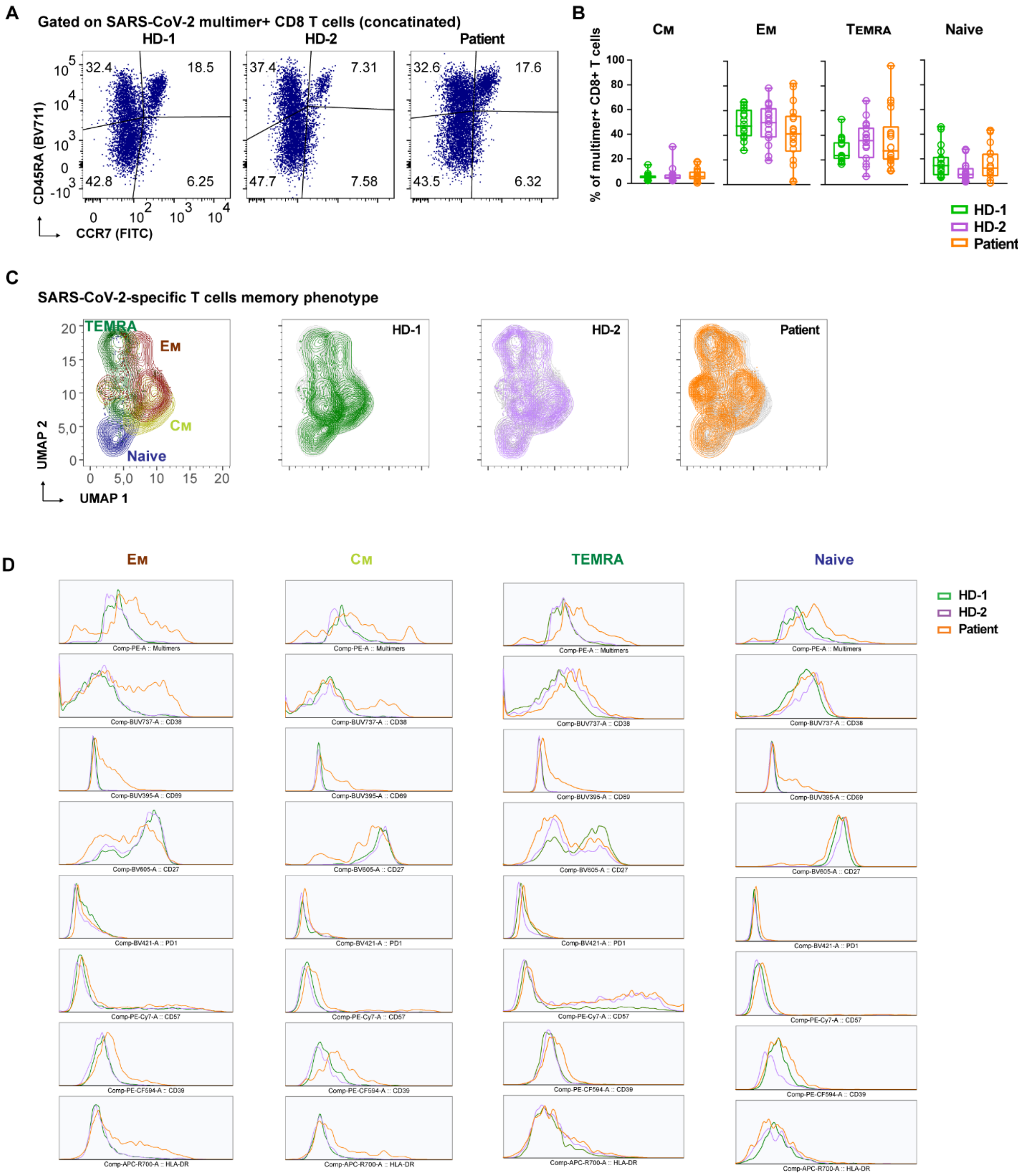


**Supplementary Figure 7. Memory phenotype of SARS-CoV-2 reactive CD8$^+$ T cells. (A)** Distribution of memory and naïve T cell phenotypes within the SARS-CoV-2 multimer$^+$ CD8$^+$ T cells identified in the patient and healthy donor cohorts based on the expression of CD45RA and CCR7 cell surface markers. Flow cytometry dot plots show the fraction of multimer$^+$ CD8$^+$ T cells displaying each of the depicted phenotype markers. Population is gated based on concatenated FCS file (n = 18 for each cohort). **(B)** Quantified frequencies of memory ($C_M$, $E_M$, and $T_{EMRA}$) and naïve SARS-COV-2 multimer$^+$ CD8$^+$ T cells compared between patient and healthy donors. Each circle represents one sample. N = 18 samples in each cohort. **(C)** UMAP (leftmost plot) of combined (all three cohorts) SARS-CoV-2 multimer$^+$ T cells showing clustering of memory and naïve populations, individual UMAP plots with the distribution depicted for each cohort. **(D)** Histogram overlays comparing memory and naïve SARS-CoV-2 multimer$^+$ T cells of the patient and healthy donors in relation to additional phenotype markers: CD38, CD39, CD69, CD27, CD57, PD-1, and HLA-DR.

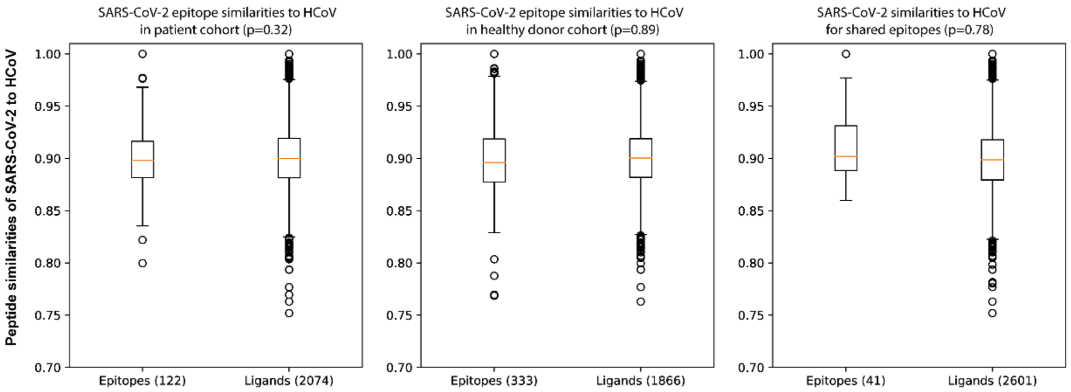


**Supplementary Figure 8. Sequence similarities of SARS-CoV-2 peptides with HCoV.** Box plots showing distribution of analyzed peptides for similarities of SARS-CoV-2-derived peptides with HCoV (-OC43, -HKU1, -NL63, and -229E) compared between epitopes (T cell reactive) and ligands (not recognizing any T cells). Data are shown for peptides analyzed in patient and healthy donor cohorts (pool of HD-1 and HD-2) and for the shared epitopes (recognized in both patient and healthy donors). P values indicate similarity values compared using welch's t-test with significance level at 0.05. The test is performed by subsampling the ligand distribution, and data analyzed using kernel method for amino acid sequences using BLOSUM62.

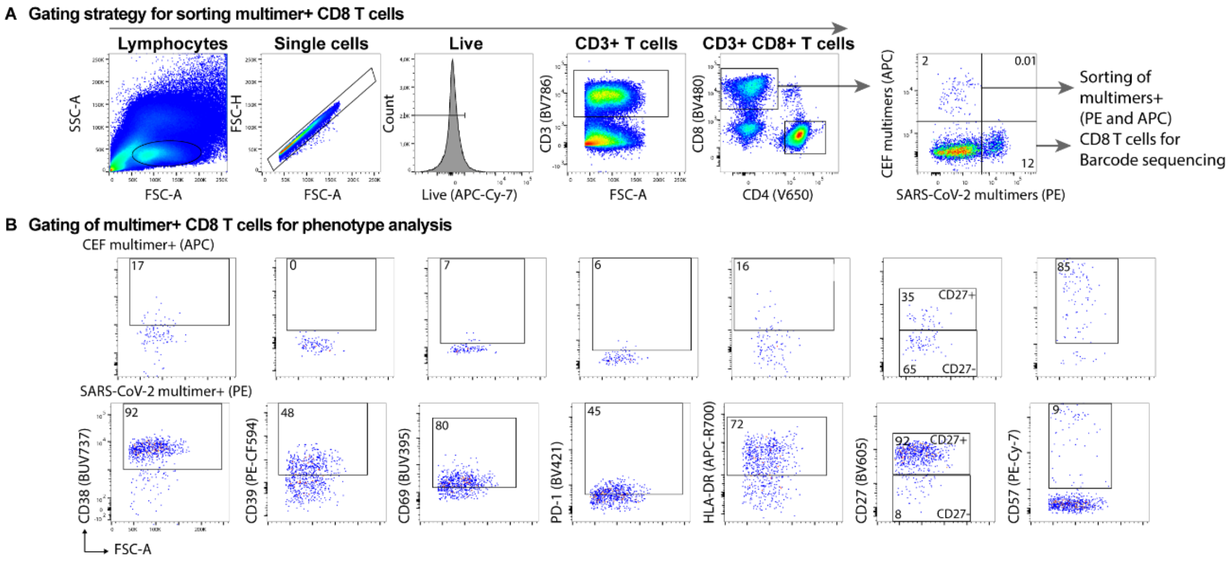


**Supplementary Figure 9. Gating strategy for sorting multimer$^+$ CD8$^+$ T cells and phenotype analysis.** Representative flow cytometry plots showing gating strategy on COVID-19 patient PBMCs stained with PE (SARS-CoV-2) and APC (CEF) conjugated DNA-barcoded pMHC multimers together with cell-surface antibodies for phenotype markers (**Supplementary Table 3**). **(A)** Gating strategy to sort SARS-CoV-2 (PE) and CEF (APC) multimer$^+$ CD8$^+$ T cells to identify individual specificities by sequencing DNA-barcodes associated with multimer$^+$ T cells. **(B)** Gating strategy applied to identify and quantify multimer$^+$ CD8$^+$ T cells expressing phenotype markers.

#### Supplementary Table 2: Patient details

| Case no. | Age | Sex | Co-morbidity no. | Medication no. | Symptoms | Days admitted no. | Days at ICU no. | Severity score: | Treatment * during admission | Hemoglobulin mmol/L | Leucocyte E9/L | Trombocyte E9/L | LDH** U/L | CRP*** mg/L | Ferrithin µg/L | Viral load# | Clinical outcome |
|---|---|---|---|---|---|---|---|---|---|---|---|---|---|---|---|---|---|
| 1 | 82 | F | 4 (heart, diabetes, stroke, endocrine) | 11 | gastro intestinal | 3 | 0 | 2 | AB | 8.5 | 6.6 | 198 | 219 | 13 | 799 | | Recovered |
| 2 | 55 | M | 1 (lymphoma) | 0 | fever, upper respiratory, lower respiratory | 13 | 0 | 3 | O2, AB, Ig | 5.9 | 1.4 | 31 | 426 | 40 | 4830 | 34.2 | Recovered |
| 3 | 48 | M | 2 (diabetes, obesity) | 2 | fever, gastro-intestinal, lower respiratory, other | 6 | 0 | 2 | O2, trial | 8.7 | 4.8 | 190 | 375 | 70 | 2020 | 16.2 | Recovered |
| 4 | 66 | M | 2 (heart, lung) | 6 | fever, lower respiratory | 30 | 21 | 4 | O2, AB | 5.4 | 9 | 393 | 334 | 56 | 1050 | | Recovered |
| 5 | 58 | M | 3 (kidney, solid cancer, autoimmunity) | 0 | fever, lower respiratory | 36 | 21 | 4 | O2, AB, pred, Ig | 5.6 | 8 | 329 | 270 | 11 | 995 | 24.4 | Recovered |
| 6 | 34 | M | 2 (diabetes, high cholesterol) | 4 | fever, gastro-intestinal, lower respiratory | 5 | 0 | 2 | O2, AB, trial | 10.8 | 8 | 203 | 306 | 119 | 948 | 13.3 | Recovered |
| 7 | 50 | F | 1 (endocrine) | 2 | Upper respiratory, lower respiratory, gastro-intestinal, other | 4 | 0 | 2 | O2, AB, trial | 7.7 | 6.1 | 207 | 342 | 35 | 277 | 31.2 | Recovered |
| 8 | 34 | F | 1 (obesity) | 0 | lower respiratory, gastro-intestinal, other | 12 | 5 | 4 | O2, AB, trial, pred | 8.1 | 3.1 | 161 | 297 | 38 | 410 | 26.5 | Recovered |
| 9 | 38 | M | 0 | 0 | fever, lower respiratory, gastro-intenstinal, other | 6 | 0 | 2 | O2, AB | 7.5 | 10 | 335 | 486 | 38 | 3960 | 25.2 | Recovered |
| 10 | 52 | M | 1 (obesity) | 0 | fever, lower respiratory, gastro-intenstinal, other | 9 | 0 | 3 | O2, AB | 7.6 | 7.8 | 212 | 327 | 66 | NA | 26.7 | Recovered |
| 11 | 29 | M | 0 | 0 | | 0 | 0 | 1 | | NA | NA | NA | NA | NA | NA | 29.1 | Recovered |
| 12 | 36 | M | 0 | 0 | | 0 | 0 | 1 | | NA | NA | NA | NA | NA | NA | 23.5 | Recovered |
| 13 | 56 | M | 1 (heart) | 2 | | 0 | 0 | 1 | | NA | NA | NA | NA | NA | NA | 17.6 | Recovered |
| 14 | 42 | M | 0 | 0 | | 0 | 0 | 1 | | NA | NA | NA | NA | NA | NA | 21.5 | Recovered |
| 15 | 41 | F | 1 (lung) | 0 | | 0 | 0 | 1 | | NA | NA | NA | NA | NA | NA | 39.7 | Recovered |
| 16 | 43 | F | 0 | 0 | | 0 | 0 | 1 | | NA | NA | NA | NA | NA | NA | 14 | Recovered |
| 17 | 44 | M | 1 (epilepsia) | 4 | lower respiratory, gastro-intestinal | 3 | 0 | 2 | O2, AB, trial | 9.1 | 6.1 | 215 | 196 | 4 | 363 | 26.6 | Recovered |
| 18 | 41 | F | 0 | 0 | | 0 | 0 | 1 | | NA | NA | NA | NA | NA | NA | 23.7 | Recovered |

*O2=oxygen, AB = antibiotic, pred = prednisolon, Ig= Immunoglobulin

** lactate dehydrogenase

***C-reactive protein

### viral load

NA-not available

Severity score: 1 -not admitted 2: admitted mild 3: admitted severe 4: severe need for ICU

**Supplementary Table 3: Phenotype antibody panel**

| Antibody | Conjugate | Clone | Dilution | Provider | Catalogue ID |
|---|---|---|---|---|---|
| CD3 | BV786 | SK7 | 1/20 | BD Biosciences | 563800 |
| CD4 | BV650 | SK3 | 1/40 | BD Biosciences | 563875 |
| CD8 | BV480 | RPA-T8 | 1/50 | BD Biosciences | 566121 |
| CD45RA | BV711 | HI100 | 1/40 | BD Biosciences | 563733 |
| CCR7 | FITC | G043H7 | 1/20 | Biolegend | 352116 |
| CD27 | BV605 | O323 | 1/40 | Biolegend | 302830 |
| CD38 | BUV737 | HB7 | 1/160 | BD Biosciences | 612824 |
| CD39 | PE-CF594 | Tu66 | 1/40 | BD Biosciences | 563678 |
| CD57 | PECy7 | QA17A04 | 1/40 | Biolegend | 393310 |
| CD69 | BUV395 | FN50 | 1/20 | BD Biosciences | 564364 |
| CD137 | PE-Cy5 | 4B4-1 | 1/40 | BD Biosciences | 551137 |
| HLA-DR | APC-R700 | G46-6 | 1/160 | BD Biosciences | 565127 |
| PD1 | BV421 | EH12.2H7 | 1/33 | BioLegend | 329920 |
| Live-Dead marker | APC-Cy7 | - | 1/1000 | Thermo Fishcer Scientific | L34976 |

#### Supplementary Table 4: CEF peptide library

| S. No. | HLA type | Peptide | Peptide sequence |
|---|---|---|---|
| 1 | HLA-A*01:01 | CMV pp65 YSE | YSEHPTFTSQY |
| 2 | HLA-A*01:01 | CMV pp50 VTE | VTEHDTLLY |
| 3 | HLA-A*01:01 | FLU BP-VSD | VSDGGPNLY |
| 4 | HLA-A*02:01 | FLU MP 58-66 GIL | GILGFVFTL |
| 5 | HLA-A*02:01 | EBV LMP2 CLG | CLGGLLTMV |
| 6 | HLA-A*02:01 | EBV BMF1 GLC | GLCTLVAML |
| 7 | HLA-A*02:01 | EBV LMP2 FLY | FLYALALLL |
| 8 | HLA-A*02:01 | CMV pp65 NLV | NLVPMVATV |
| 9 | HLA-A*02:01 | EBV BRLF1 YVL | YVLDHLIVV |
| 10 | HLA-A*02:01 | CMV IE1 VLE | VLEETSVML |
| 11 | HLA-A*02:01 | HIV Pol (C20) | ILKEPVHGV |
| 12 | HLA-A*03:01 | CMV pp150 TTV | TTVYPPSSTAK |
| 13 | HLA-A*03:01 | FLU NP 265-273 ILR | ILRGSVAHK |
| 14 | HLA-A*03:01 | EBV EBNA 3a RLR | RLRAEAQVK |
| 15 | HLA-A*03:01 | CMV pp150 TVY | TVYPPSSTAK |
| 16 | HLA-A*03:01 | EBV BRLF1 148-56 RVR | RVRAYTYSK |
| 17 | HLA-A*11:01 | EBV-EBNA4 | AVFDRKSDAK |
| 18 | HLA-A*11:01 | HCMV pp65 | GPISGHVLK |
| 19 | HLA-A*24:02 | EBV EBNA-3 114-121 | RYSIFFDY |
| 20 | HLA-A*24:02 | EBV LMP-2 419-427 | TYGPVFMCL |
| 21 | HLA-A*24:02 | EBV Rta 28-37 | DYCNVLNKEF |
| 22 | HLA-A*24:02 | HCMV 248-256 | AYAQKIFKIL |
| 23 | HLA-B*07:02 | CMV pp65 TPR | TPRVTGGGAM |
| 24 | HLA-B*07:02 | CMV pp65 RPH-L | RPHERNGFTV |
| 25 | HLA-B*07:02 | EBV EBNA RPP | RPPIFIRLL |
| 26 | HLA-B*08:01 | Flu NP (C8) | ELRSRYWAI |
| 27 | HLA-B*08:01 | EBV BZLF1 (C9) | RAKFKQLL |
| 28 | HLA-B*08:01 | CMV IE1 | ELRRKMMYM |
| 29 | HLA-B*08:01 | EBV EBNA 3A (C10) | QAKWRLQTL |
| 30 | HLA-B*08:01 | EBV EBNA 3A (C11) | FLRGRAYGL |
| 31 | HLA-B*15:01 | EBV Nuclear antigen EBNA-3C | QNGALAINTF |
| 32 | HLA-B*15:01 | EBV EBNA3A nuclear protein | LEKARGSTY |
| 33 | HLA-B*15:01 | Influenza polymerase PB1 | TQIQTRRSF |
| 34 | HLA-B*15:01 | Influenza polymerase PB1 | KMARLGKGY |
| 35 | HLA-B*15:01 | EBV EBNA3B (EBNA4A) latent protein | GQGGSPTAM |
| 36 | HLA-B*35:01 | Flu Matrix (C15) | ASCMGLIY |
| 37 | HLA-B*35:01 | EBV EBNA 3B (C16) | AVLLHEESM |
| 38 | HLA-B*35:01 | EBV BZLF1 (C17) | APENAYQAY |
| 39 | HLA-B*35:01 | CMVpp65 | IPSINVHHY |

#### Supplementary Table 5: SARS-CoV-2 -specific T cell epitopes identified in COVID-19 patients.

| HLA | Peptide | Protein | log_fold_change | p | Est. frequency | Samples analyzed |
|---|---|---|---|---|---|---|
| A01:01 | AAISDYDYY | ORF1 | 4.190 | 3.27E-13 | 0.046 | 5 |
| | CTDDNALAYY | ORF1 | 3.381 | 9.50E-07 | 0.022 | 5 |
| | CTEIDPKLDNY | ORF1 | 2.332 | 2.22E-04 | 0.050 | 5 |
| | CTEIDPKLDNY | ORF1 | 3.186 | 9.87E-08 | 0.049 | 5 |
| | FTCASEYTGNY | ORF1 | 3.486 | 2.18E-04 | 0.010 | 5 |
| | FTSDYYQLY | ORF3 | 5.683 | 1.31E-26 | 0.175 | 5 |
| | FTSDYYQLY | ORF3 | 2.166 | 6.76E-04 | 0.082 | 5 |
| | HTTDPSFLGRY | ORF1 | 5.086 | 1.36E-15 | 0.030 | 5 |
| | LGDVRETMSY | ORF1 | 2.441 | 8.99E-05 | 0.050 | 5 |
| | LLTDEMIAQY | S | 3.254 | 5.70E-06 | 0.019 | 5 |
| | PTDNYITTY | ORF1 | 6.253 | 7.32E-28 | 0.072 | 5 |
| | SNSGSDVLY | ORF1 | 2.729 | 1.17E-05 | 0.035 | 5 |
| | TDNYITTY | ORF1 | 4.441 | 2.49E-13 | 0.035 | 5 |
| | TEIDPKLDNYY | ORF1 | 4.841 | 2.65E-17 | 0.051 | 5 |
| | TIEVNSFSGY | ORF1 | 5.395 | 7.99E-25 | 0.233 | 5 |
| | TITQMNLKY | ORF1 | 6.436 | 1.79E-33 | 0.376 | 5 |
| | TSSGDATTAY | ORF1 | 4.018 | 1.04E-09 | 0.025 | 5 |
| | TTDPSFLGRY | ORF1 | 12.770 | 1.37E-88 | 11.207 | 5 |
| | TTDPSFLGRY | ORF1 | 3.558 | 1.00E-04 | 5.083 | 5 |
| | TTDPSFLGRY | ORF1 | 4.512 | 4.29E-17 | 0.196 | 5 |
| | TTDPSFLGRY | ORF1 | 5.015 | 2.66E-20 | 0.149 | 5 |
| | TTDPSFLGRY | ORF1 | 2.924 | 1.54E-06 | 0.042 | 5 |
| | TTDPSFLGRYM | ORF1 | 11.992 | 7.49E-83 | 14.109 | 5 |
| | TTDPSFLGRYM | ORF1 | 3.190 | 9.69E-09 | 0.170 | 5 |
| | TTDPSFLGRYM | ORF1 | 3.488 | 3.93E-10 | 0.111 | 5 |
| | VATSRTLSYY | M | 8.226 | 3.49E-47 | 0.350 | 5 |
| | VDDPCPIHFY | ORF8 | 6.985 | 8.20E-37 | 0.212 | 5 |
| | YFTSDYYQLY | ORF3 | 5.190 | 3.08E-17 | 0.035 | 5 |
| A02:01 | AADLDDFSKQL | N | 2.101 | 6.38E-04 | 0.007 | 8 |
| | ALSKGVHFV | ORF3 | 2.406 | 6.65E-05 | 0.055 | 8 |
| | ELDERIDKV | ORF1 | 2.389 | 8.79E-05 | 0.007 | 8 |
| | FIAGLIAIV | S | 2.676 | 1.86E-05 | 0.005 | 8 |
| | FLAHIQWMV | ORF1 | 2.764 | 1.89E-05 | 0.004 | 8 |
| | FLARGIVFM | ORF1 | 2.165 | 4.37E-04 | 0.007 | 8 |
| | FLGIITTV | ORF8 | 3.082 | 1.91E-08 | 0.031 | 8 |
| | FLPFFSNV | S | 4.627 | 1.25E-18 | 0.089 | 8 |
| | HLMSFPQSA | S | 2.732 | 1.40E-06 | 0.015 | 8 |
| | KLNVGDYFV | ORF1 | 2.860 | 3.31E-07 | 0.017 | 8 |
| | KLSYGIATV | ORF1 | 2.891 | 2.55E-07 | 0.018 | 8 |
| | KLVNKFLAL | ORF1 | 2.561 | 7.43E-06 | 0.017 | 8 |
| | KNLSDRVVFV | ORF1 | 2.499 | 7.80E-06 | 0.031 | 8 |
| | KQGDDYVYL | ORF1 | 3.137 | 1.91E-08 | 0.021 | 8 |
| | LFLLPSLATV | ORF1 | 2.427 | 2.34E-05 | 0.014 | 8 |
| | NLLLLFVTV | ORF3 | 3.574 | 5.01E-08 | 0.049 | 8 |
| | NLLLLFVTV | ORF3 | 2.736 | 5.08E-06 | 0.037 | 8 |
| | NLYDKLVSSFL | ORF1 | 2.133 | 8.89E-04 | 0.005 | 8 |
| | NPLLYDANYFL | ORF3 | 3.178 | 8.98E-08 | 0.030 | 8 |
| | RTIKVFTTV | ORF1 | 4.290 | 5.06E-13 | 0.080 | 8 |
| | RTIKVFTTV | ORF1 | 2.048 | 8.79E-04 | 0.006 | 8 |
| | SALWEIQQVV | ORF1 | 2.486 | 6.65E-05 | 0.028 | 8 |
| | SFELLHAPATV | S | 2.269 | 1.49E-04 | 0.009 | 8 |
| | SLDTYPSLETI | ORF1 | 3.095 | 2.72E-08 | 0.017 | 8 |
| | SLDTYPSLETI | ORF1 | 2.462 | 4.11E-05 | 0.015 | 8 |
| | SLLSVLLSM | ORF1 | 2.758 | 5.66E-04 | 0.032 | 8 |
| | VLNDILSRL | S | 2.884 | 3.31E-07 | 0.014 | 8 |
| | VVFLHVTYV | S | 2.397 | 3.36E-05 | 0.012 | 8 |
| | YLYFIKGLNNL | ORF1 | 2.460 | 3.40E-05 | 0.008 | 8 |
| | YMHHMELPTGV | ORF1 | 2.555 | 1.01E-04 | 0.004 | 8 |
| A03:01 | KCYGVSPTK | S | 4.916 | 2.11E-16 | 0.013 | 8 |
| | KLFDRYFKY | ORF1 | 6.284 | 6.35E-31 | 0.092 | 8 |
| | KTFPPTEPK | N | 4.404 | 2.69E-12 | 0.009 | 8 |
| A24:02 | AYANRNRFL | M | 2.202 | 5.44E-04 | 0.015 | 5 |
| | DDYQGKPLEF | ORF1 | 2.720 | 2.52E-05 | 0.010 | 5 |
| | DYKHYTPSF | ORF1 | 2.203 | 4.16E-04 | 0.022 | 5 |
| | DYYQLYSTQL | ORF3 | 2.711 | 3.55E-06 | 0.023 | 5 |
| | FYAYLRKHF | ORF1 | 4.714 | 3.92E-19 | 0.088 | 5 |
| | FYAYLRKHF | ORF1 | 3.188 | 3.86E-05 | 0.014 | 5 |
| | SFGPLVRKI | ORF1 | 2.370 | 4.09E-05 | 0.051 | 5 |
| | SVYYTSNPTTF | ORF1 | 3.552 | 1.52E-10 | 0.030 | 5 |
| | TFISAARQGF | ORF1 | 3.598 | 4.34E-11 | 0.041 | 5 |
| | TFTYASALW | ORF1 | 3.339 | 5.76E-06 | 0.054 | 5 |
| | TQDLFLPFF | S | 2.339 | 5.01E-04 | 0.009 | 5 |
| | TYVPAQEKNFT | S | 3.422 | 2.83E-06 | 0.060 | 5 |
| | VYIGDPAQL | ORF1 | 7.957 | 4.50E-46 | 0.793 | 5 |
| | VYMPASWVM | ORF1 | 2.670 | 6.78E-06 | 0.020 | 5 |
| | YLPYPDPSRIL | ORF1 | 7.005 | 5.75E-39 | 1.488 | 5 |
| | YYPSARIVY | ORF1 | 3.032 | 5.54E-08 | 0.032 | 5 |
| | YYQLYSTQL | ORF3 | 3.210 | 7.99E-09 | 0.030 | 5 |
| | YYTSNPTTF | ORF1 | 7.487 | 2.24E-42 | 0.699 | 5 |
| | YYTSNPTTFHL | ORF1 | 5.770 | 1.41E-27 | 0.179 | 5 |
| B07:02 | LEIPRRNVATL | ORF1 | 4.375 | 3.18E-16 | 0.103 | 5 |
| | MIAQYTSAL | S | 5.165 | 1.22E-22 | 0.230 | 5 |
| | SPRWYFYYL | N | 10.316 | 1.00E-67 | 8.674 | 5 |
| | SPRWYFYYL | N | 3.286 | 1.24E-07 | 0.124 | 5 |
| | SPRWYFYYL | N | 6.379 | 6.58E-33 | 0.055 | 5 |

| | | | | | | |
|---|---|---|---|---|---|---|
| | **SPRWYFYYL** | N | 3.775 | 4.38E-12 | 0.023 | 5 |
| **B08:01** | **HPLADNKFAL** | ORF7 | 3.545 | 2.53E-09 | 0.018 | 4 |
| | **LPQGFSAL** | S | 3.180 | 4.74E-07 | 0.011 | 4 |
| | **NINLHTQVV** | ORF1 | 2.392 | 9.66E-04 | 0.038 | 4 |
| | **NLKTLLSL** | ORF1 | 2.589 | 3.84E-04 | 0.006 | 4 |
| | **NRFLYIIKL** | M | 2.882 | 9.93E-06 | 0.010 | 4 |
| | **QSASKIITL** | ORF3 | 3.292 | 9.01E-08 | 0.058 | 4 |
| | **SFNPETNIL** | M | 2.508 | 1.37E-04 | 0.011 | 4 |
| | **SLYVNKHAF** | ORF1 | 3.106 | 4.94E-05 | 0.005 | 4 |
| | **YAYLRKHF** | ORF1 | 2.618 | 1.21E-04 | 0.009 | 4 |
| | **YFVVKRHTF** | ORF1 | 2.289 | 1.00E-03 | 0.049 | 4 |
| **B15:01** | **ALRKVPTDNY** | ORF1 | 2.267 | 3.68E-04 | 0.013 | 3 |
| | **CVADYSVLY** | S | 5.507 | 1.20E-25 | 0.101 | 3 |
| | **FAYANRNRF** | M | 2.552 | 3.59E-05 | 0.002 | 3 |
| | **FVVEVVDKY** | ORF1 | 2.281 | 7.71E-05 | 0.010 | 3 |
| | **HVGEIPVAY** | ORF1 | 2.255 | 9.65E-05 | 0.009 | 3 |
| | **KIEELFYSY** | ORF1 | 2.327 | 4.92E-05 | 0.012 | 3 |
| | **KMADQAMTQMY** | ORF1 | 2.085 | 5.31E-04 | 0.005 | 3 |
| | **KQFDTYNLW** | ORF1 | 3.306 | 2.41E-08 | 0.013 | 3 |
| | **LLKEPCSSGTY** | ORF7 | 2.382 | 3.77E-05 | 0.009 | 3 |
| | **LLNKHIDAY** | N | 3.529 | 3.51E-11 | 0.028 | 3 |
| | **LMVVIPDYNTY** | ORF1 | 2.122 | 4.35E-04 | 0.005 | 3 |
| | **LVKNKCVNF** | S | 2.446 | 2.18E-05 | 0.013 | 3 |
| | **RPQIGVVREF** | ORF1 | 2.161 | 1.99E-04 | 0.015 | 3 |
| | **RVAGDSGFAAY** | M | 5.365 | 1.27E-23 | 0.020 | 3 |
| | **TICAPLTVF** | ORF1 | 2.759 | 8.97E-04 | 0.000 | 3 |
| | **VASQSIIAY** | S | 2.407 | 4.07E-05 | 0.005 | 3 |
| | **VLNEKCSAY** | ORF1 | 2.629 | 4.17E-06 | 0.011 | 3 |
| | **VVQQLPETY** | ORF1 | 2.521 | 9.55E-05 | 0.008 | 3 |
| | **YLITPVHVM** | ORF1 | 3.012 | 1.03E-06 | 0.010 | 3 |
| | **YLITPVHVM** | ORF1 | 2.513 | 2.18E-05 | 0.005 | 3 |
| | **YLKLTDNVY** | ORF1 | 2.034 | 6.80E-04 | 0.006 | 3 |
| **C06:02** | **CRSKNPLLY** | ORF3 | 2.495 | 4.71E-04 | 0.002 | 2 |
| | **FYYVWKSYV** | ORF1 | 2.994 | 1.20E-07 | 0.021 | 2 |
| | **FYYVWKSYV** | ORF1 | 2.594 | 8.16E-05 | 0.003 | 2 |
| | **LRKHFSMMI** | ORF1 | 4.184 | 1.07E-14 | 0.011 | 2 |
| | **NSFSGYLKL** | ORF1 | 2.084 | 8.27E-04 | 0.055 | 2 |
| **C07:01** | **LRVEAFEYY** | ORF1 | 2.582 | 1.75E-04 | 0.038 | 6 |
| | **SAPPAQYEL** | ORF1 | 2.470 | 6.73E-05 | 0.028 | 6 |
| | **SRVLGLKTL** | ORF1 | 2.441 | 3.53E-04 | 0.041 | 6 |
| **C07:02** | **ARLYYDSMSY** | ORF1 | 2.258 | 5.30E-04 | 0.052 | 7 |
| | **FAIGLALYY** | ORF1 | 2.402 | 6.59E-05 | 0.068 | 7 |
| | **FYYVWKSY** | ORF1 | 2.854 | 4.16E-07 | 0.098 | 7 |
| | **FYYVWKSYV** | ORF1 | 3.144 | 7.68E-08 | 0.068 | 7 |
| | **FYYVWKSYV** | ORF1 | 2.132 | 7.70E-04 | 0.025 | 7 |
| | **IYKTPPIKDF** | S | 2.868 | 1.72E-07 | 0.284 | 7 |
| | **LRIMASLVL** | ORF1 | 6.595 | 1.24E-33 | 0.220 | 7 |
| | **LTDEMIAQY** | S | 2.235 | 2.83E-04 | 0.059 | 7 |
| | **LVKPSFYVY** | E | 2.627 | 3.96E-05 | 0.042 | 7 |
| | **LYLQYIRKL** | ORF1 | 2.166 | 6.27E-04 | 0.064 | 7 |
| | **MFVKHKHAF** | ORF1 | 2.098 | 7.42E-04 | 0.075 | 7 |
| | **RFDNPVLPF** | S | 3.122 | 8.51E-09 | 0.212 | 7 |
| | **SLRPDTRYVL** | ORF1 | 5.400 | 4.13E-25 | 0.356 | 7 |
| | **SLRPDTRYVL** | ORF1 | 2.222 | 5.82E-04 | 0.020 | 7 |
| | **VRIIMRLWL** | ORF3 | 2.619 | 1.55E-05 | 0.042 | 7 |
| | **YFTSDYYQL** | ORF3 | 2.155 | 6.79E-04 | 0.058 | 7 |
| | **YRYNLPTM** | ORF1 | 2.285 | 7.79E-04 | 0.034 | 7 |

**Supplementary Table 7: HLA genotype data for patient and healthy donors.**

| S. NO. | Sample ID | HLA-A | | HLA-B | | HLA-C | |
|---|---|---|---|---|---|---|---|
| **HLA genotype: Patient cohort** | | | | | | | |
| 1 | AP-02-01 | 02:01:01G | 24:02:01G | 07:02:01G | 51:01:01G | 07:02:01G | 15:02:01G |
| 2 | AP-03-01 | 01:01:01G | 24:02:01G | 08:01:01G | 15:01:01G | 07:01:01G | 07:04:01G |
| 3 | AP-04-01 | 01:01 | 24:02:01G | 39:06:02G | 49:01:01G | 07:01:01G | 07:02:01G |
| 4 | AP-05-01 | 01:01:01G | 02:01:01G | 07:02:01G | 40:01:01G | 03:04:01G | 07:02:01G |
| 5 | AP-06-01 | 03:01:01G | 24:02:01G | 08:01:01G | 44:02:01G | 07:01:01G | 07:04:01G |
| 6 | AP-08-01 | 01:01:01G | 68:01:02G | 52:01:01G | 57:01:01G | 06:02:01G | 12:02:01G |
| 7 | AP-09-01 | 03:01:01G | 11:01:01G | 40:06:01G | 55:01:01G | 01:02:01G | 15:02:01G |
| 8 | AP-10-01 | 01:01:01G | - | 08:01:01G | 57:01:01G | 06:02:01G | 07:01:01G |
| 9 | AP-11-01 | 02:01:01G | 02:05:01G | 15:01:01G | 58:01:01G | 07:01:01G | - |
| 10 | AP-12-01 | 03:01:01G | 24:02:01G | 07:02:01G | 49:01:01G | 07:01:01G | 07:02:01G |
| 11 | AP-13-01 | 02:01:01G | 32:01:01G | 27:05:02G | 44:02:01G | 02:02:02G | 07:04:01G |
| 12 | AP-14-01 | 02:01:01G | 30:02:01G | 18:01:01G | 39:01:01G | 05:01:01G | 07:02:01G |
| 13 | AP-15-01 | 02:01:01G | 03:01:01G | 08:01:01G | 40:01:01G | 03:04:01G | 07:01:01G |
| 14 | AP-16-01 | 02:01:01G | 03:01:01G | 35:01:01G | 44:02:01G | 04:01:01G | 05:01:01G |
| 15 | AP-17-01 | 03:01:01G | 30:01:01G | 15:01:01G | - | 03:03:01G | 03:04:01G |
| 16 | AP-18-01 | 03:01:01G | 31:01:02G | 07:02:01G | 27:05:02G | 01:02:01G | 07:02:01G |
| 17 | AP-19-01 | 02:01:01G | 30:01:01G | 15:01:01G | 44:02:01G | 03:03:01G | 05:01:01G |
| 18 | AP-20-01 | 02:01:01G | 03:01:01G | 07:02:01G | 44:02:01G | 05:01:01G | 07:02:01G |
| **HLA genotype: HD-1 cohort** | | | | | | | |
| 1 | BC-10 | 02:01:01G | | 07:02:01G | 15:01:01G | 04:01:01G | 07:02:01G |
| 2 | BC-55 | 01:01:01G | 02:01:01G | 08:01:01G | 40:01:01G | 07:01:01G | 03:04:01G |
| 3 | BC-67 | 02:01:01G | 30:01:01G | 07:02:01G | 13:02:01G | 06:02:01G | 07:02:01G |
| 4 | BC-80 | 01:01:01G | 02:01:01G | 08:01:01G | 57:01:01G | 06:02:01G | 07:01:01G |
| 5 | BC-100 | 01:01:01G | 02:01:01G | 08:01:01G | 15:01:01G | 07:01:01G | 03:04:01G |
| 6 | BC-103 | 11:01:01G | 24:02:01G | 35:01:01G | 40:01:01G | 04:01:01G | 03:04:01G |
| 7 | BC-111 | 02:01:01G | 69:01:01 | 15:01:01G | 55:01:01G | 03:03:01G | 03:04:01G |
| 8 | BC-120 | 01:01:01G | 02:01:01G | 08:01:01G | 44:02:01G | 07:01:01G | 05:01:01G |
| 9 | BC-329 | 02:01:01G | 25:01 | 07:02:01G | 35:01:01G | 04:01:01G | 07:02:01G |
| 10 | BC-335 | 02:01:01G | | 35:01:01G | 40:01:01G | 04:01:01G | 03:04:01G |
| 11 | BC-336 | 01:01:01G | 24:02:01G | 08:01:01G | 44:03:01G | 07:01:01G | 16:01:01G |
| 12 | BC-327 | 02:01:01G | 03:01:01G | 27:05:02G | 51:01:01G | 02:01:02G | 02:02:02G |
| 13 | BC-13 | 03:01:01G | 24:02:01G | 07:02:01G | 44:03:01G | 04:01:01G | 07:02:01G |
| 14 | BC-37 | 03:01:01G | | 07:02:01G | | | 07:02:01G |
| 15 | BC-69 | 01:01:01G | 03:01:01G | 08:01:01G | 35:01:01G | 04:01:01G | 07:01:01G |
| 16 | BC-71 | 02:01:01G | 03:01:01G | 07:02:01G | 44:02:01G | 05:01:01G | 07:02:01G |
| 17 | BC-84 | 03:01:01G | 24:02:01G | 07:02:01G | 35:01:01G | 04:01:01G | 07:02:01G |
| 18 | BC-85 | 01:01:01G | 03:01:01G | 07:02:01G | 08:01:01G | 07:01:01G | 07:02:01G |
| **HLA genotype: HD-2 cohort** | | | | | | | |
| 1 | BS-01-01 | 02:01:01G | 26:01:01G | 15:01:01G | 44:02:01G | 03:04:01G | 07:04:01G |
| 2 | BS-02-01 | 02:01:01G | 24:02:01G | 14:02:01G | 15:01:01G | 03:04:01G | 08:02:01G |
| 3 | BS-04-01 | 02:01:01G | 03:01:01G | 07:02:01G | 52:01:01G | 07:01:01G | 07:02:01G |
| 4 | BS-06-01 | 01:01:01G | 03:01:01G | 07:02:01G | 27:05:02G | 02:02:02G | 07:02:01G |
| 5 | BS-09-01 | 02:01:01G | 03:01:01G | 07:02:01G | 13:02:01G | 06:02:01G | 07:02:01G |
| 6 | BS-11-01 | 01:01:01G | 02:01:01G | 08:01:01G | 18:01:01G | 07:01:01G | - |
| 7 | BS-12-01 | 01:01:01G | - | 08:01:01G | 27:05:02G | 01:02:01G | 07:01:01G |
| 8 | BS-13-01 | 03:01:01G | 24:02:01G | 07:02:01G | 44:03:01G | 07:02:01G | 16:01:01G |
| 9 | BS-14-01 | 02:01:01G | 24:02:01G | 15:01:01G | 40:02:01G | 02:02:02G | 03:04:01G |
| 10 | BS-16-01 | 02:01:01G | 02:02:01G | 07:02:01G | 53:01:01G | 04:01:01G | 07:02:01G |
| 11 | BS-17-01 | 02:01:01G | 03:01:01G | 07:02:01G | - | 07:02:01G | - |
| 12 | BS-18-01 | 03:01:01G | 26:01:01G | 07:02:01G | 44:03:01G | 04:01:01G | 07:02:01G |
| 13 | BS-19-01 | 11:01:01G | 24:02:01G | 55:01:01G | 57:01:01G | 03:03:01G | 06:02:01G |
| 14 | BS-20-01 | 01:01:01G | 29:02:01G | 08:01:01G | 44:03:01G | 07:01:01G | 16:01:01G |
| 15 | BS-23-01 | 02:01:01G | 03:01:01G | 07:02:01G | 15:01:01G | 03:03:01G | 07:02:01G |
| 16 | BS-30-01 | 01:01:01G | 02:01:01G | 08:01:01G | 51:01:01G | 03:03:01G | 07:01:01G |
| 17 | BS-35-01 | 01:01:01G | 02:01:01G | 07:02:01G | 08:01:01G | 07:01:01G | 07:02:01G |
| 18 | BS-37-01 | 01:01:01G | 24:02:01G | 08:01:01G | 18:01:01G | 07:01:01G | - |
| 19 | BS-38-01 | 01:01:01G | 03:01:01G | 13:02:01G | 40:02:01G | 02:02:02G | 06:02:01G |
| 20 | BS-39-01 | 01:01:01G | 24:02:01G | 07:02:01G | 08:01:01G | 07:01:01G | 07:02:01G |
